# Antibody Upstream Sequence Diversity and Its Biological Implications Revealed by Repertoire Sequencing

**DOI:** 10.1101/2020.09.02.280396

**Authors:** Yan Zhu, Xiujia Yang, Jiaqi Wu, Haipei Tang, Qilong Wang, Junjie Guan, Wenxi Xie, Sen Chen, Yuan Chen, Minhui Wang, Chunhong Lan, Lai Wei, Caijun Sun, Zhenhai Zhang

**Affiliations:** State Key Laboratory of Organ Failure Research, National Clinical Research Center for Kidney Disease, Division of Nephrology, Nanfang Hospital, Southern Medical University, Guangzhou, 510515, China; Department of Bioinformatics, School of Basic Medical Sciences, Southern Medical University, Guangzhou 510515, China; Center for Precision Medicine, Guangdong Provincial People’s Hospital, Guangdong Academy of Medical Sciences, Guangzhou 510080, China; Key Laboratory of Mental Health of the Ministry of Education, Guangdong-Hong Kong-Macao Greater Bay Area Center for Brain Science and Brain-Inspired Intelligence, Southern Medical University, Guangzhou 510515, China; Department of Nephrology, Hainan Affiliated Hospital of Hainan Medical College, Haikou 570311, China; Department of Nephrology, Hainan General Hospital, Haikou 570311, China; State Key Laboratory of Ophthalmology, Zhongshan Ophthalmic Center, Sun Yat-sen University, Guangzhou 510060, China; School of public health, Sun Yat-sen University, Shenzhen 510006, China

**Keywords:** antibody upstream sequences, 5’ UTR, leader sequences, Rep-Seq, antibody repertoire

## Abstract

The sequence upstream of antibody variable region (Antibody Upstream Sequence, or AUS) consists of 5’ untranslated region (5’ UTR) and two leader regions, L-PART1 and L-PART2. The sequence variations in AUS affect the efficiency of PCR amplification, mRNA translation, and subsequent PCR-based antibody quantification as well as antibody engineering. Despite their importance, the diversity of AUSs has long been neglected. Utilizing the rapid amplification of cDNA ends (5’RACE) and high-throughput antibody repertoire sequencing (Rep-Seq) technique, we acquired full-length AUSs for human, rhesus macaque (RM), cynomolgus macaque (CM), mouse, and rat. We designed a bioinformatics pipeline and discovered 2,957 unique AUSs, corresponding to 2,786 and 1,159 unique sequences for 5’ UTR and leader, respectively. Comparing with the leader records in the international ImMunoGeneTics (IMGT), while 529 were identical, 313 were with single nucleotide polymorphisms (SNPs), 280 were totally new, and 37 updated the incomplete records. The diversity of AUSs’ impact on related antibody biology was also probed. Taken together, our findings would facilitate Rep-Seq primer design for capturing antibodies comprehensively and efficiently as well as provide a valuable resource for antibody engineering and the studies of antibody at the molecular level.

## Introduction

Antibodies represent an essential class molecule constituting adaptive immune system and can specifically bind to the invading pathogens for the subsequent eradication or clearance [1]. Rep-Seq technology has enabled antibodies to be interrogated in an unprecedented coverage and depth, by which researchers can obtain millions to even billions of antibody sequences in a single experiment [2-4]. The application of Rep-Seq has led to a substantial progress in many fields, such as aging, tumor immunology, infectious diseases, immune surveillance and neutralizing antibody screening [5,6].

Albeit to the successes mentioned above, the potentials of Rep-Seq can be confined by a series of not fully addressed issues both experimentally and computationally [5]. One of the primary computational roadblocks is the unknown germline sequence diversity [7-9]. The uncaptured diversity will lead to germline misassignment and thus bias the downstream analyses. Besides, the germline polymorphisms of immunoglobulin loci are found to associate with expressed antibody repertoire and disease predisposition [10-13]. To capture these diversity, many tools were developed and employed in antibody repertoire studies [7-9], which then led to the findings of many novel alleles.

Despite the progresses mentioned above, the diversity of AUSs were less interrogated. An AUS contains two consecutive functional elements, namely 5’ UTR and leader, both of which were found to implicate in mRNA transcription and translation [14-21]. Particularly, leader sequences play a vital role in antibody expression and are often engineered to improve the efficiency of monoclonal antibody production [16,21,22]. In Rep-Seq studies, the AUSs are often the targets of the PCR primers for obtaining full-length antibody variable regions [23,24]. Furthermore, Mikocziova et al. showed that polymorphisms in AUSs can facilitate the annotation of antibody variable (V) genes [25]. Owing to their importance aforementioned, we decided to interrogate the possible diversity of AUSs which were long neglected in the field.

We therefore sequenced the antibody repertoire of both heavy and light chains of 5 species, namely human, rhesus macaque (RM), cynomolgus macaque (CM), mouse and rat. Applying the devised bioinformatic pipeline to the in-house dataset together with available public resources, we discovered thousands of unique AUSs. We then examined their functional relevance to the expression and status of downstream V genes within as well as among species. Our findings here provided the first overview of antibody AUS features of multiple species and enriched the relevant knowledge database which would serve as valuable resources for the community.

## Results

### Bioinformatics pipeline for obtaining high-quality AUSs

As shown in Figure 1, we obtained the candidate AUSs through the following six steps: i) V allele variant detection via IgDiscover with initial species-specific databases downloaded from IMGT/GENE-DB [7,26]. The newly discovered genes/alleles were merged with initial databases and carried on for downstream analyses; ii) the sequences from each sample were annotated and assembled via MiXCR (compared in Zhang et al. [27]) and clonotypes were consequently extracted based on CDR3s and V and junctional (J) allele usages; iii) each sample were genotyped via a Bayesian method adapted from TIgGER; iv) the AUS in each clonotype was extracted and dimed as the initial AUS for that particular V allele; v) for each V allele, the consensus of all initial AUSs were calculated and defined as the final AUS(s). Alleles not in the genotype were excluded in this step; vi) a scoring-based method was employed to retain the most confident AUSs. Further details on each of these steps can be found in Materials and Methods.

**Figure 1.**
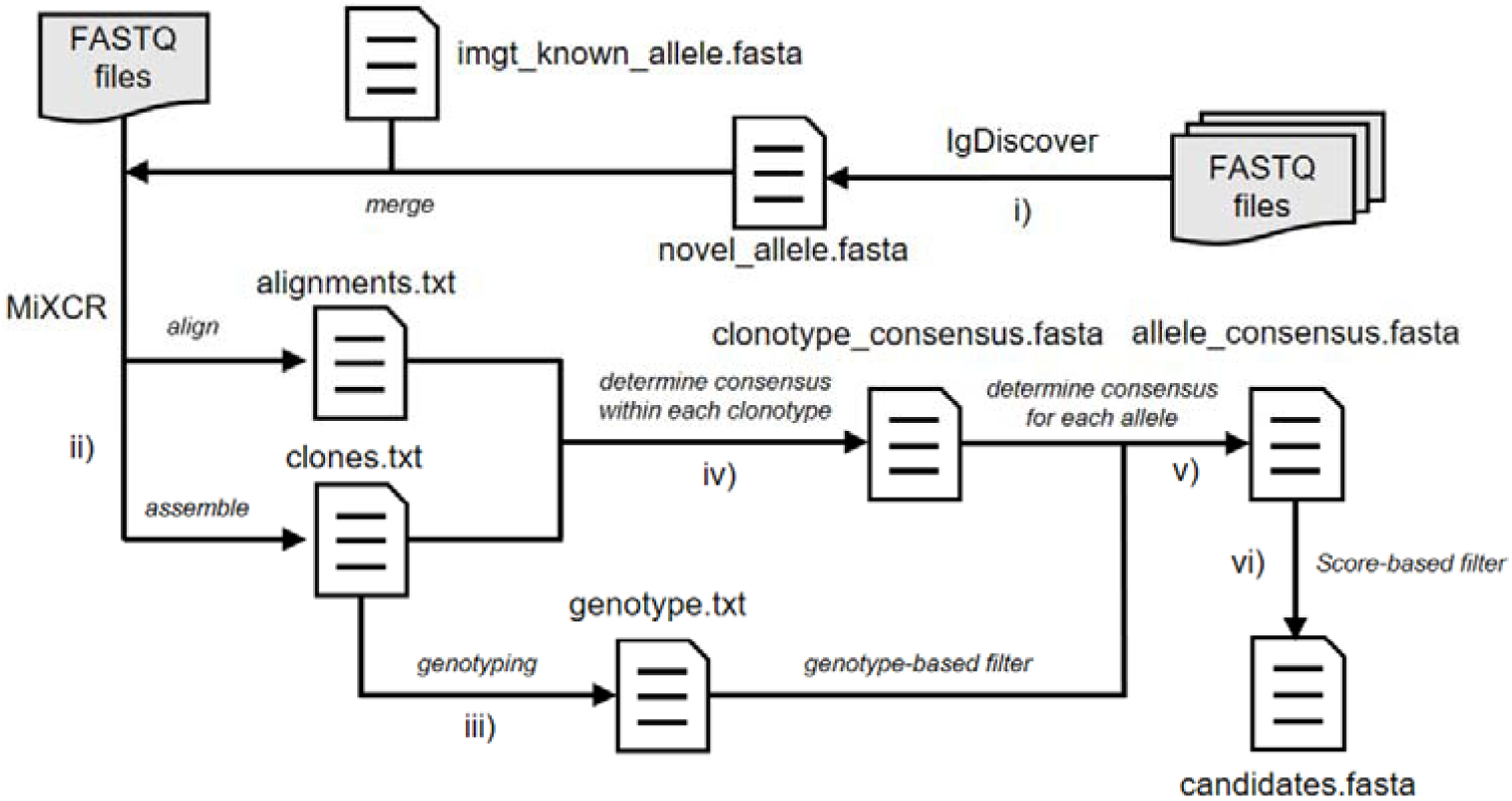
The AUS identification pipeline. The AUS identification pipeline is comprised with six steps, including **i)** V allele variant detection, **ii)** sequence annotation and assembly, **iii)** genotyping, **iv)** identification of consensus sequences within clonotypes, **v)** identification of consensus sequences within V alleles, and **vi)** score-based filtration.

### Leaders are more conserved than 5’ UTRs

Utilizing the bioinformatics pipeline introduced above, we found 2,957 unique AUSs, corresponding to 2,786 and 1,159 unique sequences for 5’ UTR and leader in all five species, respectively (Table 1 and Supplementary Table 1). The percentage of V genes with discovered AUSs ranged from 7.7% (52 genes for kappa chain of RM) to 56.9% (33 genes for lambda chain of human). For all chain types across five species, we discovered AUSs for up to 73.9% of the genes with complete leaders in IMGT/GENE-DB. Of these, 70% human heavy chain AUSs displayed previously unreported polymorphisms (Supplementary figure 1 and Supplementary Table 2) which may affect Rep-Seq capture efficiency and antibody production. For human V gene alleles of unknown or incomplete leader according to IMGT/GENE-DB, 43, 26, and 26 (47.3%, 56.5% and 55.3%) AUSs were revealed for heavy, kappa and lambda chain V genes, respectively.

**Table 1.** Overview of discovered AUSs.

| Species | Chain | # unique AUSs | # unique 5' UTRs | # unique leaders | # alleles | # genes | % genes |
| --- | --- | --- | --- | --- | --- | --- | --- |
| Human | IGH | 1111 | 1023 | 346 | 205 | 61 | 47.3 |
|  | IGk | 498 | 458 | 189 | 138 | 43 | 44.3 |
|  | IGL | 161 | 152 | 76 | 65 | 33 | 56.9 |
| RM | IGH | 5 | 5 | 2 | 2 | 2 | 3.8 |
|  | IGK | 50 | 48 | 31 | 24 | 24 | 20.3 |
| CM | IGH | 136 | 132 | 69 | 52 | 52 | 35.1 |
| Mouse | IGH | 239 | 237 | 118 | 110 | 106 | 30.8 |
|  | IGK | 256 | 256 | 96 | 93 | 91 | 54.2 |
| Rat | IGH | 399 | 379 | 162 | 126 | 126 | 32.3 |
|  | IGK | 102 | 96 | 70 | 60 | 60 | 36.8 |
| **Total** | **-** | **2957** | **2786** | **1159** | **-** | **-** | **-** |

Besides, we found the largest number of unique novel sequences for both leader and 5’ UTR in human, probably due to the largest sample size for human among studied species (Figure 2a and Supplementary figure 2a). This demonstrated high diversity of AUSs in human that should not be neglected and the power of analyzing large dataset. We observed the highest ratio of leader sequences consistent with known ones in mouse and rat but the lowest in RM. It is worth mentioning that the 5’ UTR sequences curated by IMGT are either partial or contain additional lengthy intron and upstream regulatory elements [25]. Therefore, a considerable number of 5’ UTRs were part of their counterparts on IMGT or *vice versa*. For CM, all 132 5’ UTRs discovered in this study were novel.

**Figure 2.**
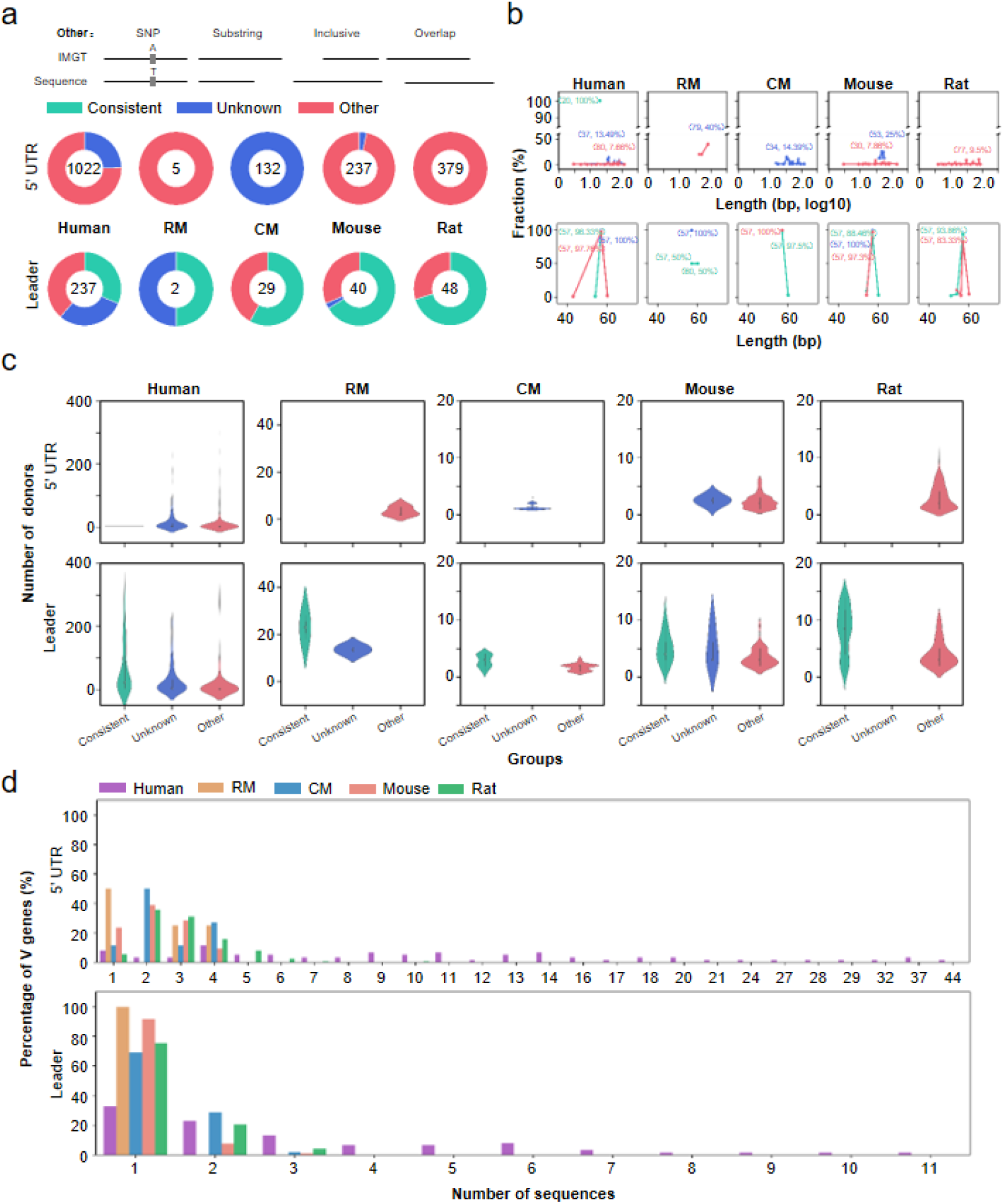
Overview of the discovered AUSs for heavy chain. **(a)** The composition of discovered 5’ UTR and leader sequences. “Consistent” indicates sequences identical to those curated by IMGT/GENE DB. “Unknown” indicates sequences whose corresponding alleles were not provided with available AUSs. “Other” includes the rest four situations, which were illustrated on top of the donut charts. The number of novel sequences is marked in the center of each donut chart. **(b)** The length distribution of discovered 5’ UTR and leader sequences. The length and its corresponding frequency were marked for each peak. **(c)** The distribution of the number of donors sharing a certain sequence. Different colors in (b) and (c) mean different groups and are consistent with those in (a). **(d)** Percentage of V genes as a function of the number of discovered 5’ UTR and leader sequences.

For all species, more than half of the leaders for heavy and kappa were 57 bp in length (Figure 2b and Supplementary figure 2b). In contrast, the lambda chain exhibited a mode length of 60 bp. The length distribution of 5’ UTRs were characterized by long tails (for example, ranging from 3 to 133 bp for human heavy chain) and peaked at different lengths for different species. Thus, the conservation of the leader sequence length across species might be a result of its functional importance. Population wise, we found the leaders consistent with IMGT records showed similar frequencies in donors with those different from IMGT, which indicated the reliability of the newly identified AUSs (Figure 2c and Supplementary figure 2c). Individual genes exhibited high diversity with regard to their AUSs. Only less than 5% and around 30% of human V genes used single unique 5’ UTRs and leaders, respectively. A single V gene may use up to 44 different 5’ UTRs (human IGHV1-69) and 11 leader sequences (human IGHV3-53) (Figure 2d). This diversity was consistent in all five species (Supplementary figure 3). The higher diversity of 5’ UTR compared to leader was also reflected in the combinatorial frequencies, in which 54 out of 57 (94.7%) genes have more diverse 5’ UTR than leader (Supplementary figure 4), and the more consistent result of leader than 5’ UTR with Mikocziova et al. [25] (Supplementary figure 5). Taken together, these results indicated that leader sequences are more conservative than 5’ UTRs’ and more critical for the functionality of antibodies.

### The AUSs coevolved with V genes

We further investigated the sequence-level similarities among AUSs within and across species (Materials and Methods). Within the same species, we found both 5’ UTR and leaders showed clear family-specific sequence feature for both heavy and light chains (Figure 3a, b and Supplementary figure 6, 7). For human heavy chains, leader sequence sharing was only observed for genes in the same family (Figure 3c, d and Supplementary figure 8). It is also the case for 5’ UTR sequences for human light chains (Supplementary figure 8). However, a single 5’ UTR sequence (ACC) was shared between VH1 (IGHV1-3, IGHV1-17 and IGHV1-69) and VH3 (IGHV3-13) families (Figure 3c). This family wise sharing of 5’ UTRs also exists in nonhuman species (Supplementary figure 9). Thus the AUSs, in general, are V gene family specific. Besides, these within-family interchangeable leaders would also affect the application of elevating V gene assignment accuracy by incorporating upstream sequences.

**Figure 3.**
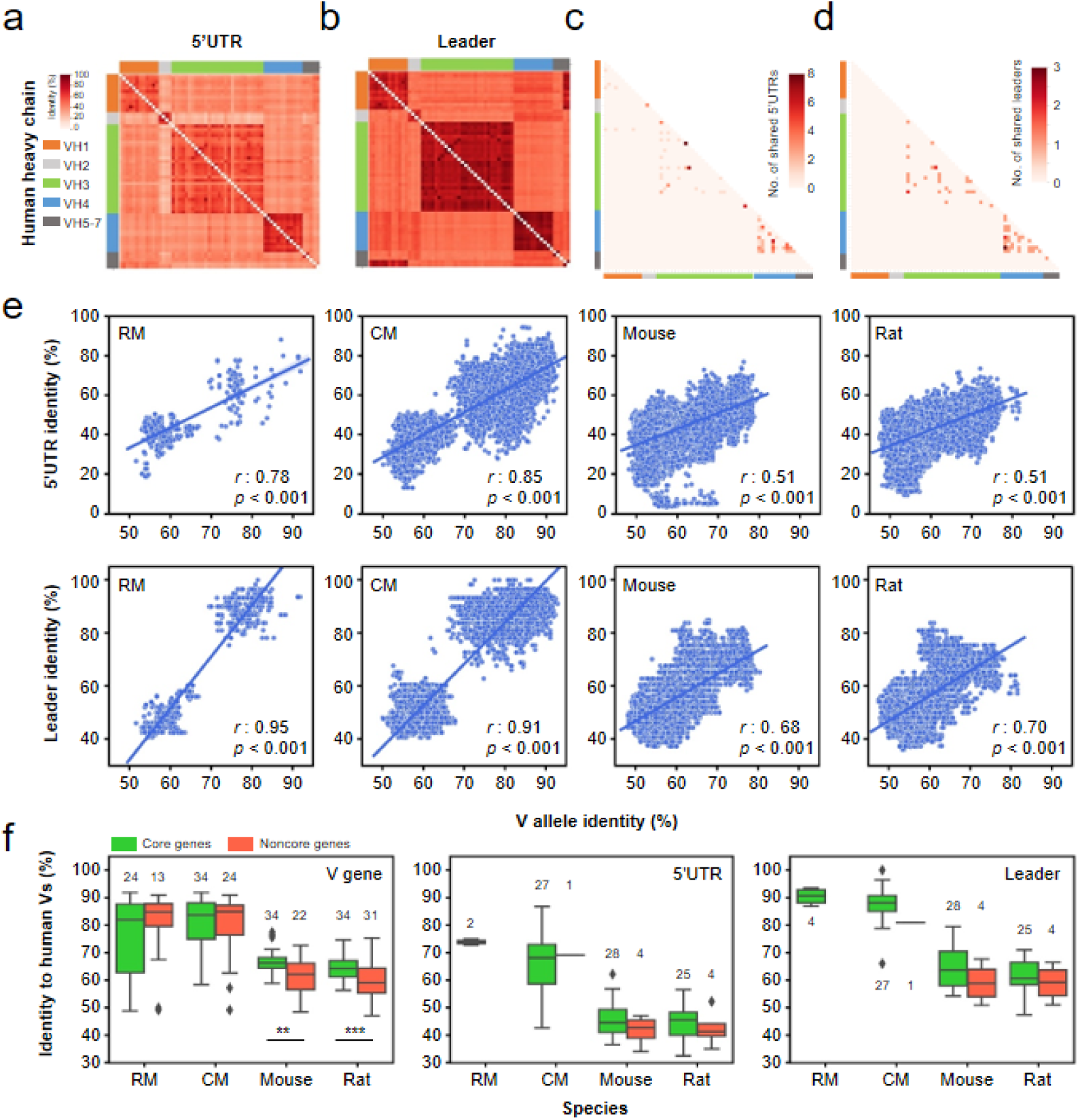
5’UTR and leader sequence similarity within human and between species. **(a)** 5’UTR sequence identity between V genes. **(b)** Leader sequence identity between V genes. **(c)** The number of shared 5’UTR sequences between V genes of different families. **(d)** The number of shared leader sequences between V genes of different families. The side color bar indicates gene families. **(e)** The correlation between 5’UTR or leader sequence identities and V allele sequence identities between human and four nonhuman species. The line in each scatterplot represents the fitted linear regression model. **(f)** V gene, 5’UTR and leader sequence identity between human and nonhuman species for antibody heavy chain. Genes were classified into core genes and noncore genes according to Yang et al., 2019. *r*, Pearson’s correlation coefficient; *p*, p value of the linear model; **, *p*<0.01; ***, *p*<0.001.

For each of the nonhuman V alleles, we identified its human counterpart by sequence similarity (Materials and Methods). Then we compared the similarities of nonhuman V gene elements to those of their human counterparts. The pairwise similarities for both 5’ UTRs and leaders correlate with V allele similarities. This indicate the AUSs evolved together with V gene alleles (Figure 3e and Supplementary figure 10). Moreover, the correlations of leaders were higher than those of 5’ UTRs. This result suggested that AUSs coevolved with V genes and the leaders and V genes were under more similar selective pressure. In details, we found that 5’ UTR and leader sequences of RM (73.9% and 90.6% for heavy chain for 5’ UTR and leader, respectively) and CM (68.4% and 87.8%) were more similar to human than mouse (44.5% and 62.2%) and rat (45.2% and 60.6%) (Figure 3f and Supplementary figure 11, Materials and Methods). This was consistent with the genomic distances reported before [28-31]. We previously classified heavy chain V genes into two groups, namely core genes and noncore genes^31^, where the former contributes to the vast majority of antibody repertoire and the latter is negligible. We then classified the heavy chain V genes for four nonhuman species and calculated the sequence similarities between nonhuman and human for V genes, 5’ UTRs, and leaders. Although noncore genes are more functionally inactive, they showed no significant differences in sequence similarity between human and nonhuman species, except for the V genes of mouse and rat (Figure 3f). This result indicated that core genes and noncore genes are of equal importance evolutionarily, despite their difference in contribution to antibody repertoire. Furthermore, we observed a roughly equal similarity between leaders and V genes, implicating again a coevolution between them.

### Leader SNPs are position-dependent and may contribute to functional reversal and inefficient amplification of V genes

We next examined the single nucleotide polymorphisms (SNPs) in the AUSs as they were reported to affect antibody production [16] and the primer design for Rep-Seq and genetic predisposition screening [13,24]. Applying a heuristic algorithm, we investigated the ratio of replacement (R) to silent (S) SNPs and positional nucleotide and amino acid diversity index (NDI and ADI) of discovered leader sequences (Materials and Methods) except for RM due to the low number of SNPs. For heavy chain leaders, the R/S ratio of human was the lowest (0.43), which was followed by CM (0.63), mouse (2.66) and rat (4.05) (Figure 4a). Similar decreased R/S ratios were also observed in light chains (Supplementary figure 12). Knowing that each leader sequence contains a hydrophobic central region [16], we classified the amino acids according to their polarities and looked into the amino acid conversion types for replacement SNPs. It demonstrated that intra-group conversions dominated overall amino acid conversions (Figure 4b and Supplementary figure 12). And the percentage of inter-group conversions followed the same order as that of replacement SNPs in different species. Furthermore, both NDI and ADI were calculated to evaluate the positional nucleotide and amino acid diversity. The NDI profile exhibited distinct patterns (Figure 4c). To be specific, the third nucleotides in 2nd, 3rd, 9th, 12th and 19th codons in leader sequences possessed clearly higher NDI than the first two nucleotides in the same codons. Since the choice of the third nucleotides in a codon often do not change the encoded amino acids, the higher NDI of the third nucleotides demonstrated a clearly negative selection over the polymorphisms taking place in the first two nucleotides in these codons. In contrast, the 13th, 14th, 16th, and 17th codons were observed with higher NDI in the first two nucleotides, suggesting the underlying positive selection. While for the 6th and 7th codons, the diversities were comparable among the constituent nucleotides. The positional ADI profile was consistent with NDI (Figure 4c).

**Figure 4.**
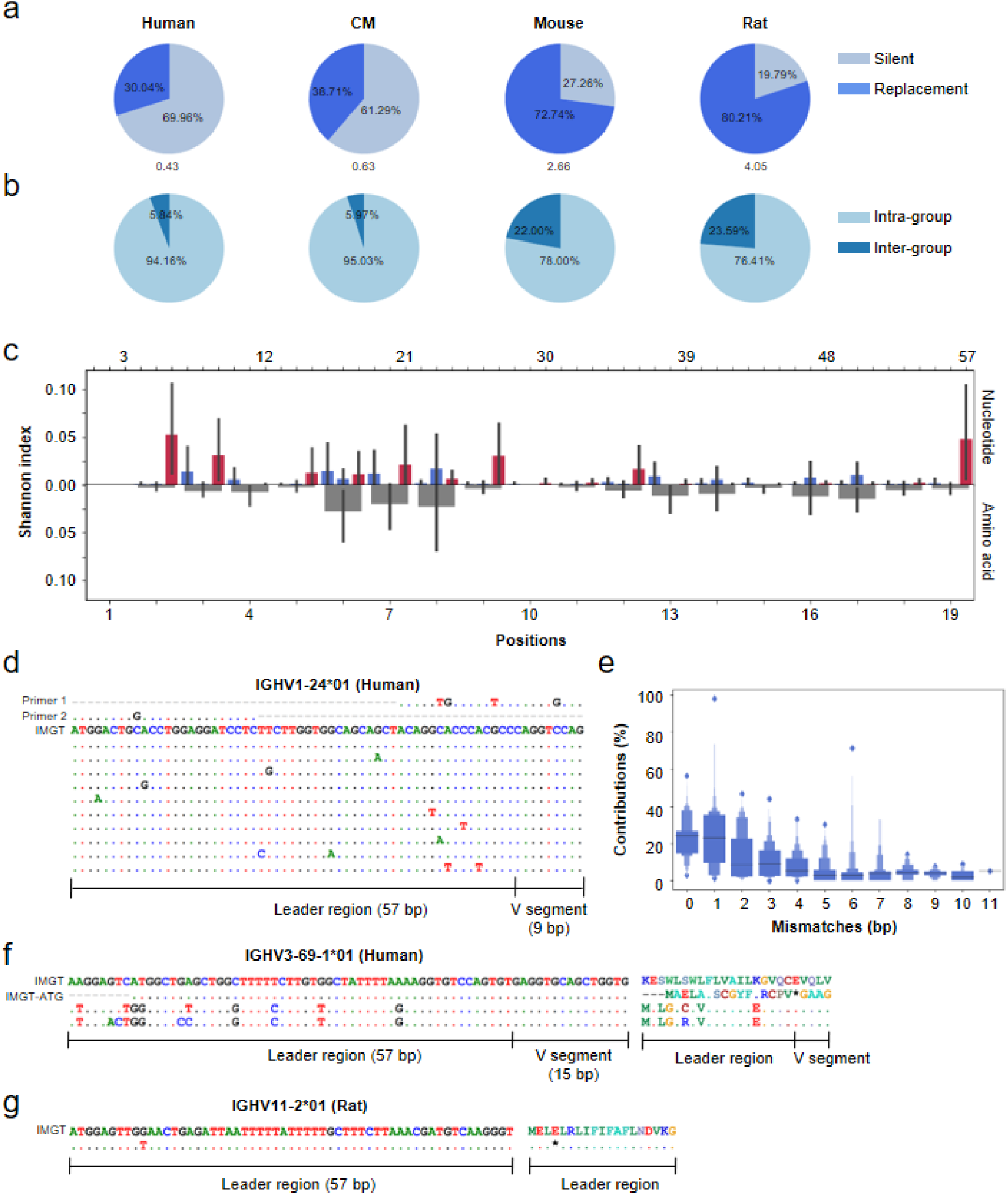
Characterization of SNPs observed in discovered leader sequences. **(a)** The percentage of silent (S) and replacement (R) SNPs for heavy chain. R/S ratio was marked beneath each pie plot. **(b)** The percentage of intra-group and inter-group amino acid conversion. Note that amino acids here were classified according to their polarities. **(c)** Positional nucleotide diversity profile of leader sequences. The vertical lines mean the standard errors. Boxes in red represent the third nucleotides in codons. **(d)** The schematic representation of mismatches of published primers with discovered leader sequences. **(e)** The correlation between the contribution to the overall target sequences and the number of their mismatches with primers. **(f, g)** Schematic representation of SNP-associated V gene functional reversals, including pseudo-to-functional (f) and functional-to-pseudo transitions (g).

The SNPs in AUSs would also affect the antibody genotyping as well as Rep-Seq when these polymorphic regions were targeted by PCR primer design. We selected two representative primer sequences from public resources that target either the 3’ end (Primer 1) or 5’ end (Primer 2) of leader sequences of IGHV1-24*01. We found both of them covered the SNP loci (5 for primer 1 and 2 for primer 2) (Figure 4d). To comprehensively evaluate the influence of novel leaders on antibody sequence amplification, we compared the novel leader sequences to 6 widely applied primer sets [24,32-36]. As a result, 9 to 22 genes were found with novel sequences more distant to the optimal primers than their counterpart in IMGT (Supplementary Table 3). Particularly, the number of mismatches can increase from 4 to 9 for IGHV1-17 in the primer set reported by Klein et al. [32] (Supplementary Table 4). Indeed, when comparing amplifications of V alleles between 5’RACE and multiplex primers, the more mismatches in the primer, the less V allele were found in the repertoire (Figure 4e, Materials and Methods). More mismatches between primers and their targeted AUSs led to more compromised amplification. Therefore, the AUS set we reported here would be a valuable resource for later primer designs.

More importantly, the SNPs in the leader region also caused functional resurrection or silence to their corresponding genes. For instance, the human IGHV3-69-1*01 and IGHV3-69-1*02 were both annotated as pseudo genes at IMGT because of the missing initiation codons at the starts of leader sequences. However, we identified 2 (supported by 34 and 5 subjects) and 1 (3 subjects) novel leader sequences that possessed normal initiation codon (*AUG*) and thus made these alleles functional (Figure 4f and Supplementary figure 13a). This functional recurrence also happened in IGHV1-67*01 of mouse (5 subjects) (Supplementary figure 13b). On the contrary, the SNP in the leader region caused early stop codon and silenced IGHV11-2*01 in rat (2 subjects) (Figure 4g). Thus, the SNPs in the AUSs are important for the activation of downstream V genes and additional awareness should be paid in the future.

### uAUGs and leader sequences do not affect antibody expression

Upstream open reading frames (uORFs) or upstream AUGs (uAUGs) have been reported to exist in 44%-50.5% of human annotated transcripts, implicate longer 5’ UTR sequences, and associate with significant reduction of downstream protein expression and modestly to markedly decreased mRNA level [14,18,37]. Having obtained 5’ UTR sequences from hundreds of unique AUSs for human, we investigated the prevalence of uAUGs in antibody V genes and its correlation with gene expression.

We observed uAUGs in the 5’ UTR of 12.4%, 11.85% and 8.70% of AUSs for human heavy, kappa and lambda chain, respectively (Table 2). Thus the uAUG frequency in antibody V genes is less than that in human mRNAs and the stochastic estimation, indicating the purifying selection [37,38]. Thus, this underrepresentation of uAUGs encoded in antibodies indicated that antibodies tend not to be subjected to this post-transcriptional regulation. We also observed a significant length difference between uAUG-containing and uAUG-absent 5’ UTRs (Figure 5a). Notably, these differences did not fully agree with what was reported in previous studies. The antibody heavy chain 5’ UTRs from human and CM demonstrated a reversed pattern, in which uAUG-containing 5’ UTRs were even shorter. This result implied that a longer 5’ UTR is not necessarily predisposed to contain uORFs and thus further address the polymorphism of post-transcriptional regulation for different antibody chains and for different species.

**Table 2.** Number and percentage of genes with identified uAUGs.

| **Species** | **Chain** | **# uAUG-containing**<br>**AUSs** | **% uAUG-containing**<br>**AUSs** | **# uAUGs-containing**<br>**genes** | **% uAUGs-containing**<br>**genes** |
| --- | --- | --- | --- | --- | --- |
| Human | IGH | 138 | 12.4 | 23 | 37.7 |
|  | IGK | 59 | 11.9 | 12 | 27.9 |
|  | IGL | 14 | 8.7 | 4 | 12.1 |
| RM | IGH | 0 | 0.0 | 0 | 0.0 |
|  | IGK | 32 | 64.0 | 15 | 62.5 |
| CM | IGH | 33 | 24.3 | 21 | 40.4 |
| Mouse | IGH | 75 | 31.4 | 40 | 37.7 |
|  | IGK | 69 | 27.0 | 36 | 39.6 |
| Rat | IGH | 65 | 16.3 | 27 | 21.4 |
|  | IGK | 28 | 27.5 | 18 | 30.0 |

**Figure 5.**
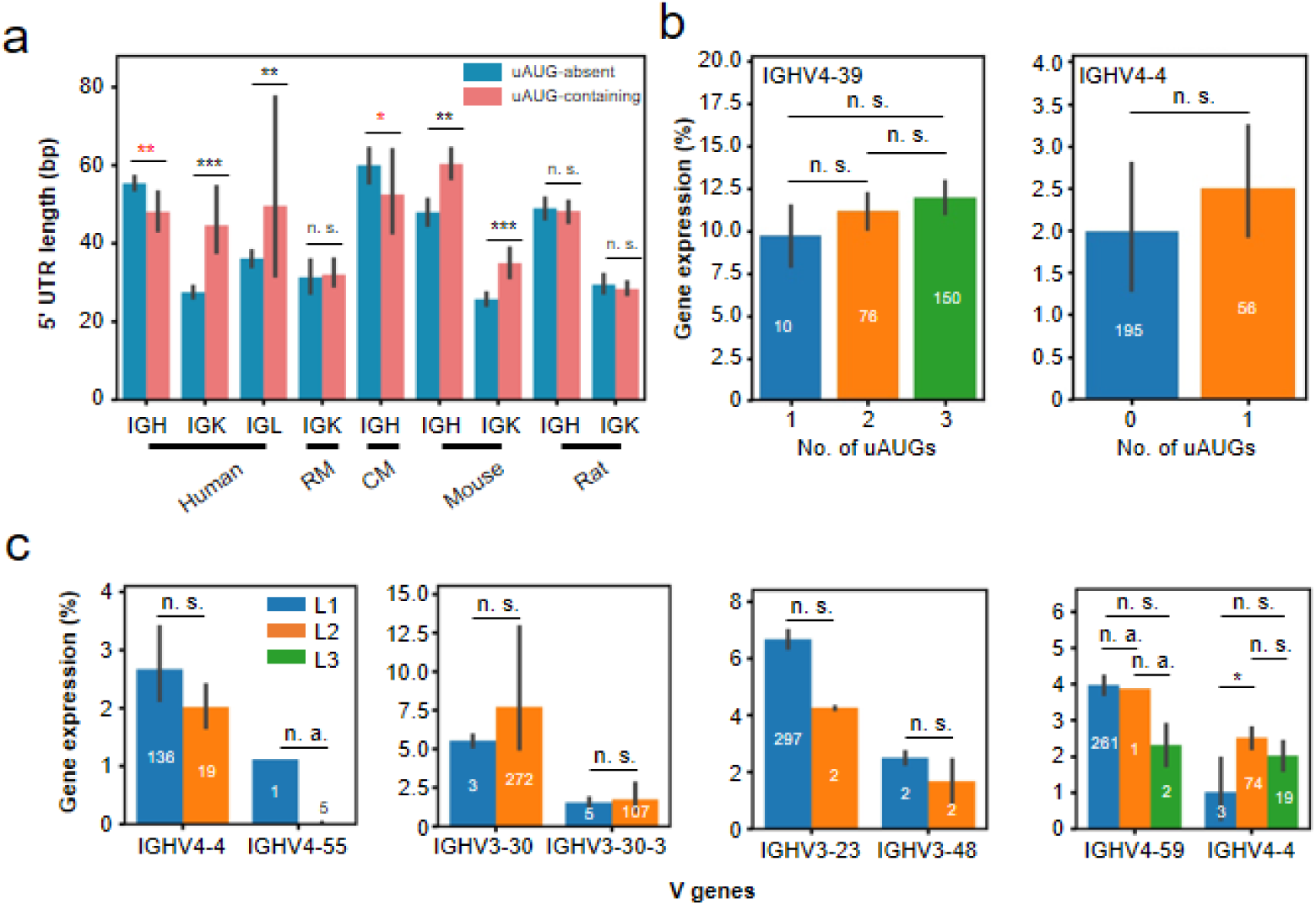
Functional relevance of uAUGs and leader sequences to gene expression. **(a)** The 5’ UTR length comparison between uAUG-containing group and uAUG-absent group in different species. The red asterisks marked the chains with reversed length difference between two groups. **(b)** The functional relevance of the number of uAUGs in 5’ UTR sequences to gene expression for human heavy chain V genes. **(c)** The functional relevance of leader sequences to gene expression for human heavy chain V genes. L1, L2, and L3 represent unique leader sequences. L1 and L2 in different subplots are not necessarily the same. The number in the center of or on top of each bar mean the number of samples in each group. The vertical lines mean the standard errors. n. a. not available; n. s., not significant (*p*>=0.5); *, *p*<0.05; **, *p*<0.01; ***, *p*<0.001.

The high throughput Rep-Seq approach gives us access to the measurement of gene expression. We thus obtained the gene expression for each sample and correlated it with the number of uAUGs present in the 5’ UTR sequences of their corresponding V genes. Two and four genes meeting the criteria were included in this analysis for human heavy and kappa chain, respectively (Materials and Methods). Although varied mRNA level was observed in different uAUG number group, statistical analysis indicated the difference was not significant (p>0.5, unpaired Student’s t-test), except for a kappa gene - IGKV1-17 (Figure 5b and Supplementary figure 14). Together with the observation that the correlation was ambiguous (both seemingly positive and negative correlation were obtained), we believed that the influence of uAUGs on antibody expression, if any, is not critical. It is noteworthy that 21 of 23 heavy chain V genes with which we observed uAUGs are core genes which are prevalent in the repertoire [39] (Supplementary figure 15). As half of the heavy chain V genes are noncore genes that express very lowly in the repertoire, the uAUGs in these genes might be unrevealed in this study.

Apart from uORFs, the leader sequence itself was also found to associate with mRNA expressions [15,20]. To validate whether and to what extent leader sequences affect antibody transcription, we set off to investigate the antibody expression of V genes sharing two or more unique leaders. As shown in Figure 5c and Supplementary figure 16, the overall pattern showed no significant leader-specific gene expression was found. Thus the leader sequences involve more in the regulation of translation rather than transcription. Altogether, the results above reflected the polymorphism of antibody 5’ UTR sequences across species and chains and the trivial functional relevance to gene expression of uORFs and leader sequences.

## Discussion

Rep-Seq has provided an avenue by which researchers are able to investigate antibodies with an unprecedented depth [5]. Taking advantage of Rep-Seq, we investigated the polymorphisms existing in the AUSs of both heavy chain and light chain V genes from five species. We designed a bioinformatic pipeline and captured 2,957 unique AUSs, of which 875 (29.6%) are novel (as to leader). We demonstrated that the SNPs in the AUSs affect the PCR amplification efficiency and thus the subsequent antibody quantification. Therefore, our findings are of importance for antibody gene genotyping, antibody quantifications with traditional PCR method and high-throughput Rep-Seq. Moreover, the novel AUSs revealed in this study also provide a resource for antibody engineering and aid antibody studies in the molecular level.

In-depth analyses showed that leader sequences are family-specific and are more conserved than 5’ UTR. The distance of the AUS between human and other nonhuman species are reminiscent of that observed in genome level. And the sequence similarities between human and nonhuman species also indicates a synchronous evolution between leaders and their downstream V genes.

Notably, we found a 5’ UTR sequence was shared by V genes of different families. The length of the associated 5’ UTR was only 3 bp. It was unlikely to be the result of RNA degradation during library preparation, because the 5’cap is required for a successful template switch when using 5’RACE protocol [40]. Moreover, we found the 5’ UTR for IGHV1-17*02 in 6 subjects, a strong evidence for its authenticity. Previous studies reported a single-nucleotide 5’ UTR is enough for the initiation of translation in vitro [41]. Therefore, the short 5’ UTR may not mean the functional abnormality. However, short 5’ UTRs were found to impact the translation efficiency [41]. Due to the unavailability of proteomic data, we were not able to investigate its translation efficiency.

Informed by previous reports that uAUGs and leader sequence have an effect on gene expression in other context [4,15,18,19], we also examined if there exists correlations between them in antibody genes. We showed little evidence to support the link between them, which means for antibodies the AUSs primarily dictate the translation process. Noteworthy is that we investigated the correlations only in human, partly because we have no enough samples for nonhuman species. The other reason, however, is that there are a limited number of germline reference available [7-9]. Even worse is that some germline reference provided by IMGT/GENE-DB are provisional because their genomic locations has not been determined [26]. Hence, we proposed that the germline sequences of immunoglobulin loci should be thoroughly characterized in these species. The accurate antibody probing in these model organisms are supposed to boost the understanding of antibody repertoire in human.

In addition to the little quantitative impact on gene expression, we also observed that the SNPs in leader sequences could reverse the functional status of their downstream genes. It addressed importance of dissecting the AUSs in the evaluation of antibody functionalities, because a functional gene in a typical individual or ethnic group may be nonfunctional in another and *vice versa*. However, the prevalence of such SNP-associated functional recurrences or silences, together with their consequence for the antibody repertoire, remains to be determined. Another interesting phenomenon we observed is the higher R/S ratio in leader region for mouse and rat than human and CM. Moreover, the percentage of inter-group amino acid conversion was also higher for the two rodents. It is possibly because the immunoglobulin loci are less essential for the survival of rodents and thus they can tolerate more functional consequence caused by inter-group amino acid conversions [42].

In summary, we provided for the first time the most comprehensive knowledge database for AUSs for both human and other model organisms. Together with the characterization of these sequences, the AUS set discovered in this study will serve as valuable resources for fundamental studies and antibody engineering as well.

## Funding

This study was supported by the National Natural Science Foundation of China (NSFC) (31771479) (Z. Z.), NSFC Projects of International Cooperation and Exchanges of NSFC (61661146004), and the Local Innovative and Research Teams Project of Guangdong Pearl River Talents Program (2017BT01S131).

## Author Contributions

Y. Z., X. Y., J. W., S. C., Y. C., and C. L. analyzed the data. H. T., Q. W., J. G., W. X., and M. W. conducted the biological experiment. C. L. coordinated the project. Y. Z., X. Y., J. W., H. T., L. W., C. S., and Z. Z. wrote the manuscript. Z. Z. conceived the project.

## Competing of interests

The authors declared no competing financial interests.

## Materials and Methods

### Subjects and sample preparation

In this study, we included a total number of 782 samples from five species, including human (n=728), RM (*Macaca mulatta*) (n=27), CM (*Macaca fascicularis*) (n=2), mouse (*Mus musculus*) (n=11), and rat (*Rattus norvegicus*) (n=14). Twenty-one and twenty-six samples are from public resources for human and RM, respectively. And their accession numbers in NCBI Sequence Read Archive are provided as Supplementary Table 5. The in-house human samples were prepared as described previously [39]. As for samples from nonhuman species, EDTA-treated blood specimens were collected from sixteen animals (seven rats, six mice, two CMs, and an RM). Peripheral blood mononuclear cells (PBMCs) were isolated by Ficoll-Paque density centrifugation. The PBMCs were subjected to total RNA extraction using the RNeasy Mini Kit (Qiagen, 74106), according to the manufacturer’s protocol. This protocol was approved by the Ethics Committee of Southern Medical University.

### Library preparation and high throughput sequencing

Total RNA was used as a template to synthesize cDNA, using a SMARTer RACE (Rapid Amplification of cDNA Ends) cDNA Amplification Kit (Clontech, 634928), according to the manufacturer’s protocol. Following cDNA synthesis, 10% of the volume of cDNA was subjected to VH amplification in a 25 µl PCR reaction using the Kapa HiFi HotStart Ready Mix (KAPA Biosystems, kk2602) with universal or VH-family specific forward primers and the corresponding reverse primers (Supplementary Table 6). The thermal cycling conditions were programmed as follows: 95 °C for 3 min; 30 cycles of 98 °C for 20 s, 60 °C for 15 s, and 72 °C for 15 s; 72 °C for 5 min. PCR products were purified using the Nucleospin Gel & PCR Clean-up kit (Macherey-Nagel, 704609.25). DNA concentration was detected using the Qubit 4.0 fluorometer (ThermoFisher Scientific). Two hundred nanograms of each gel-purified product was subjected to library preparation, followed by sequencing by Illumina platform (MiSeq PE300 and NovaSeq PE250).

### Extraction of AUSs and delimitation of 5’ UTR and leader region

The Six steps we employed to identify qualified AUSs were briefly described in the Result section and demonstrated as Figure 1. Here we provided a more detailed implementation of these steps.

i. V allele variant detection. Considering the uncaptured polymorphisms in immunoglobulin loci in mammalian, especially for nonhuman species, we started with the identification of novel alleles for all enrolled samples and species [9,43,44]. We executed three iterations of IgDiscover (v0.12.3) based on an initial species-specific V gene database obtained from IMGT/GENE-DB (update: 06 July 2020) [7,26]. To enroll as many as possible the initial germline sequences, we included all sequences from the F+ORF+all P directory. The full set of germline reference was then subjected to a deduplication process before serving as the starting database. After the detection, all discovered novel alleles were merged with the initial ones for downstream sequence annotation.
ii. Sequence annotation and assembly. We then proceeded with the functionalities provided by MiXCR (v3.0.7) [45]. Taking advantage of the subcommand *align* and *assemble* and their exportation counterparts, we obtained both the sequence annotation (alignments.txt) and clustering (clones.txt) results. Sequences with the same V alleles, J alleles, and CDR3 nucleotide sequences were assembled into clonotypes. The commands we used to annotate and assemble sequences with MiXCR (v3.0.7) are as below, Alignment:

~~~
mixcr align --species $species -f --library $species.specific.library $read1 $read2
alignments.vdjca
~~~ Assembly:

~~~
mixcr assemble -f -a -OseparateByV=true -OseparateByJ=true alignments.vdjca
clones.clna
~~~ Exportation:

~~~
mixcr exportAlignments -f -readIds -cloneId -vHit -vAlignment -targetSequences
clones.clna alignments.txt
mixcr exportClones -f --chains $CHAIN clones.clna clones.txt
~~~
iii. iii) Genotyping. To guide the AUS discovery, the genotype was predetermined based on allele usage for each V gene. In this study, we considered for each V gene two scenarios, homozygous and heterozygous. Theoretically, if a gene is heterozygous in a typical sample, each of the two variants will possess a unique AUS. Otherwise, the only variant of a homozygote will probably link with two unique AUSs. We determined the genotype through the Bayesian method employed in TIgGER [46], except that we considered only two scenarios aforementioned. In this case 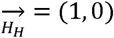 and 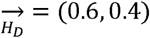. Other parameters for the proposed Bayesian model were unchanged. The allele usage, measured as the number of clonotypes recombined from a typical allele, served as the input of this function.
iv. iv) Identification of consensus sequences within clonotypes. Each AUS is deemed to start at a position immediately downstream the rGrGrGs in the 3’ end of RACE primer and end at the exclusively initial base of the associated V gene segment. For each clonotype, we extracted a list of valid AUSs and determined the most frequent one as the consensus AUS. For reliability, we discarded the consensus sequences representing no more than 50% of the AUSs within clonotypes. We also required the initiation codon – “AUG” – is contained in the AUS to precisely delimit 5’ UTR and leader region. For AU Ss with multiple AUGs, we selected an optimal AUG as the start position of leaders. Each optimal AUG corresponds to a leader sequence with its length nearest to a predetermined optimal leader length. The optimal leader length for each allele is same as the length of leader sequence provided by IMGT provided it is known. Otherwise, it is same as the most frequent leader length of all other alleles corresponding to the same gene, family or all the rest alleles with known leader sequences. Noted that we extracted only leader sequences for samples from BioProject PRJNA503527 for their UTRs were found to be incomplete. The extracted leader sequences were then subjected to the same pipeline as AUSs.
v. v) Identification of consensus sequences within V alleles. The definition of clonotype ensured that each of consensus sequences is associated with a unique V allele. AUSs corresponding to alleles not present in the genotype list were discarded at the first place. Then we collapsed the consensus sequences belonging to the same alleles and kept the most frequent ones. Alleles with no more than 10 available AUSs were excluded. For each allele in the genotype list, we retained also the second most frequent AUSs to account for the underlying diversity in the upstream region, providing the corresponding gene is a homozygote and the most frequent one takes up less than 87.5% of total AUSs [8].
vi. vi) Score-based filtration. For each chain of a typical species, the candidate AUSs for all samples were pooled together and then classified into two groups according to prior knowledge. The group containing known leader sequences were regarded as *bona fide* and thus serve as the positive control. The other group, comprised of novel sequences, will be subject to a score-based filtration. The score scheme takes into account four features for each candidate sequence, namely the absolute number of supportive reads, clones and donors and the similarity to known sequences. The weight assigned to them were 20, 20, 30, and 30, respectively. The independent scores (*S*_*read*_, *S*_*clone*_, *S*_*donor*_ and *S*_*similarity*_) and total score (*S*_*total*_) were calculated as the formulas below,

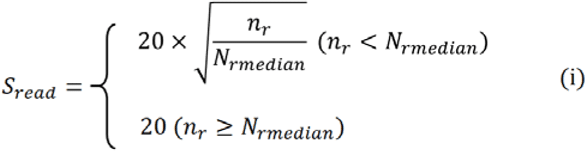

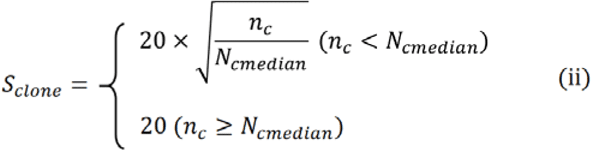

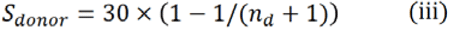

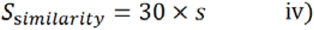

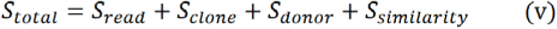

Note, *N*_*rmedian*_ and *N*_*cmedian*_ mean the median number of supportive reads and clones for known AUSs that serve as positive control while *n*_*r*_, *n*_*c*_, and *n*_*d*_ mean the number of supportive reads, clones, and donors for novel AUSs. *S* represents the identity to known leader sequences for the leader region in each AUS. If no known leader sequence was provided by IMGT/GENE-DB for the corresponding allele, an average identity to the leaders of all other alleles corresponding to the same gene or the same family or of all the rest alleles with known leader sequences was calculated to represent the identity. Thus, the theoretical maximum score for an AUS is less than 100. After the *S*_*total*_ were calculated for each AUS, the novel sequences with a score less than the median *S*_*total*_ of positive control were discarded and the rest novel sequences together with positive control serve as the final AUS library. The score-based filtration step was applied to each chain of each species independently. An exception was with the filtration of AUSs for the kappa chain of RM. Since no known complete leader sequences were discovered, the AUSs containing the known partial leader sequences were regarded as the positive control for filtration.

### The calculation of the number of unique AUS, 5’ UTR, and leader sequences

The number of unique AUS, 5’ UTR, and leader sequences is calculated as the number of unique combinations of V allele and AUS, 5’ UTR, and leader sequences. The AUSs containing only leader sequences (n=4) identified from RM samples under BioProject PRJNA503527 were not included in this analysis, but instead were included in all other analyses.

### Sequence similarity comparison

Sequence similarity or identity is calculated based on the pairwise alignment result. The pairwise alignment was implemented using *pairwise2* function built in Biopython (v1.70). After the pairwise alignment result was obtained, we calculated the sequence similarity as the number of matched nucleotides divided by the alignment length. For the measurement of leader or 5’ UTR sequence similarity between two genes within the same species, we performed an all-to-all comparison between all leader sequences corresponding to the two compared genes and calculated an average similarity for each gene pair. While for the sequence comparison between genes from human and nonhuman species, we determined the homologous genes at first. The homologous gene in human for each allele of nonhuman species was determined by the sequence similarity and the gene corresponding to the nearest allele in human was considered as its homologous gene. In this case, a typical gene in nonhuman species can have alleles with different homologous genes in human. Thus a gene in nonhuman species can be homologous with multiple genes in human and vice versa.

### Evaluation of functional relevance of uAUGs and leader sequence to gene expression

The functional relevance of uAUGs and leader sequence to gene expression was investigated in only human samples due to the limitation of sample size. To avoid the noise caused by heterozygous 5’ UTR for a gene, only genes with one unique 5’ UTR sequence were considered in each sample. Besides, among samples, only genes with diversity in the number of uAUGs and having at least 5 samples in each group were finally included in the analysis. Applying these criteria to all genes, we discovered 2 and 4 genes for heavy and light chain, respectively (Figure 5b and Supplementary figure 14). The gene filtration criteria for the evaluation of leader sequences’ influence on gene expression is similar. In each sample, only genes with one unique leader sequence were retained. Besides, among samples, only genes with at least two kinds of leader sequences and shared at least two leader sequences with another were finally included. The gene expression in each sample is calculated as the number of antibody sequences assigned a typical gene divided by the number of all assigned antibody sequences.

### The measurement of percentage of replacement and silent SNPs

Owing to the limited number of discovered leader sequence, we did not include RM into this analysis. To measure the percentage of replacement and silent SNPs, the SNP loci were firstly determined by aligning leader sequences from the same genes and then identifying the polymorphic positions. For heavy chain and lambda chain, only leader sequences of 57 bp were considered, while for kappa chain only 60 bp. This length limitation makes the leader sequence alignment quite straightforward and at the same time retain the overwhelming majority of leader diversity (Figure 2b). Then the SNP loci-associated codons were identified. For each SNP loci-associated codon position (e. g. from 1 to 19 for heavy chain), the frequencies of different codons were calculated. Each pair of codons will associate with a single SNP outcome, either replacement or silent. Then for a certain codon position, its contribution to silent SNPs was calculated as the sum of the product of the two frequencies for all codon pairs encoding the same amino acids. Similarly, its contribution to replacement SNPs was calculated as the sum of the product of the two frequencies for all codon pairs encoding different amino acids. Finally, the total replacement or silent SNPs were the sums of the contributions of all SNP loci-associated codon positions. The percentages of two kinds of amino acid conversion, namely intra-group conversion and inter-group conversion, were determined in the same way, except that it considered only replacement SNPs.

### Nucleotide and amino acid diversity index (NDI and ADI) calculation

Only SNPs of human heavy chain leader were included in this analysis. We used NDI (or ADI) to measure the nucleotide (or amino acid) diversity of each position. The diversity index was supposed to reflect the conservation of each position in the leader. Both NDI (57 positions) and ADI (19 positions) were based on the formula calculating Shannon index as follows:

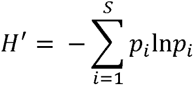

where *S* represents the number of unique nucleotides or amino acids in a certain position and *p*_i_ represents the proportion of *i*th nucleotide or amino acid.

### Mismatch comparison between known and novel leaders

In total 6 widely applied primer sets were included to compare the number of mismatches between primers and known leaders in IMGT/GENE-DB with that between primers and novel leaders found in this study. Therefore, the comparison considered only V alleles with both discovered novel leaders and their complete counterparts in IMGT/GENE-DB. For each leader sequence, we determined its optimal primer as well as the target region in each primer set by pairwise alignment. This process also enables us to obtain the number of mismatches with its optimal primer. In this way, a maximum number of mismatches with optimal primers can be obtained for each V gene (Supplementary Table 4).

### Evaluation of primer amplification efficiency

Two amplification strategies (5’RACE and multiplex PCR) were applied to the same sample and the paired sequencing dataset were acquired using NovaSeq 6000 sequencing system. We genotyped and extracted the AUSs together with the starting 30 bp sequence of V alleles for the sequenced sample using the RACE-derived dataset. For the multiplex-derived dataset, we also employed MiXCR to annotate the antibody sequences. The antibody sequences assigned to V alleles absent in the genotype were excluded at first. Then the contribution of each primer to one of its target V alleles was measured as this V allele’s percentage of all the sequences targeted by this primer. It is also straightforward to calculate the mismatches between the primer and its target V alleles based on the AUSs and partial sequences of V alleles. When correlating the contributions with mismatches, we found a clear negative correlation between them (Figure 4e).

### Software

In-house scripts were written in Python (v3.7.4) employing modules including numpy (v1.16.4), Biopython (v1.73), Levenshtein (v0.12.0) and pandas (v0.24.2). To visualize these results, we employed modules seaborn (v0.9.1) and matplotlib (v3.0.2) as well as a standalone software, BioEdit Sequence Alignment Editor.

## Notes

### Competing Interest Statement

The authors have declared no competing interest.

