## Supplementary figures for "Antibody Upstream Sequence Diversity and Its Biological Implications Revealed by Repertoire Sequencing"

### Slide 1
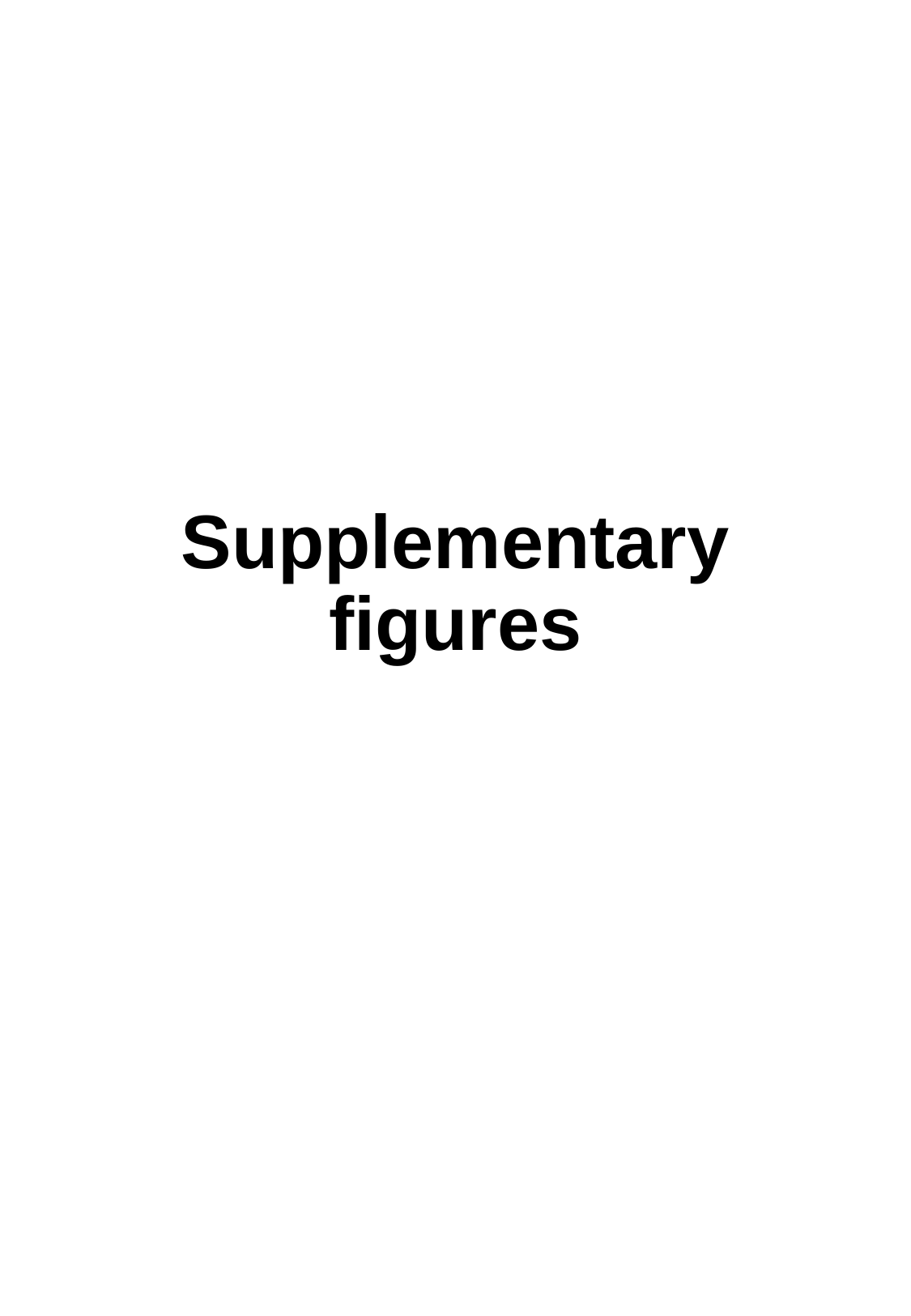

Supplementary figures

### Slide 2
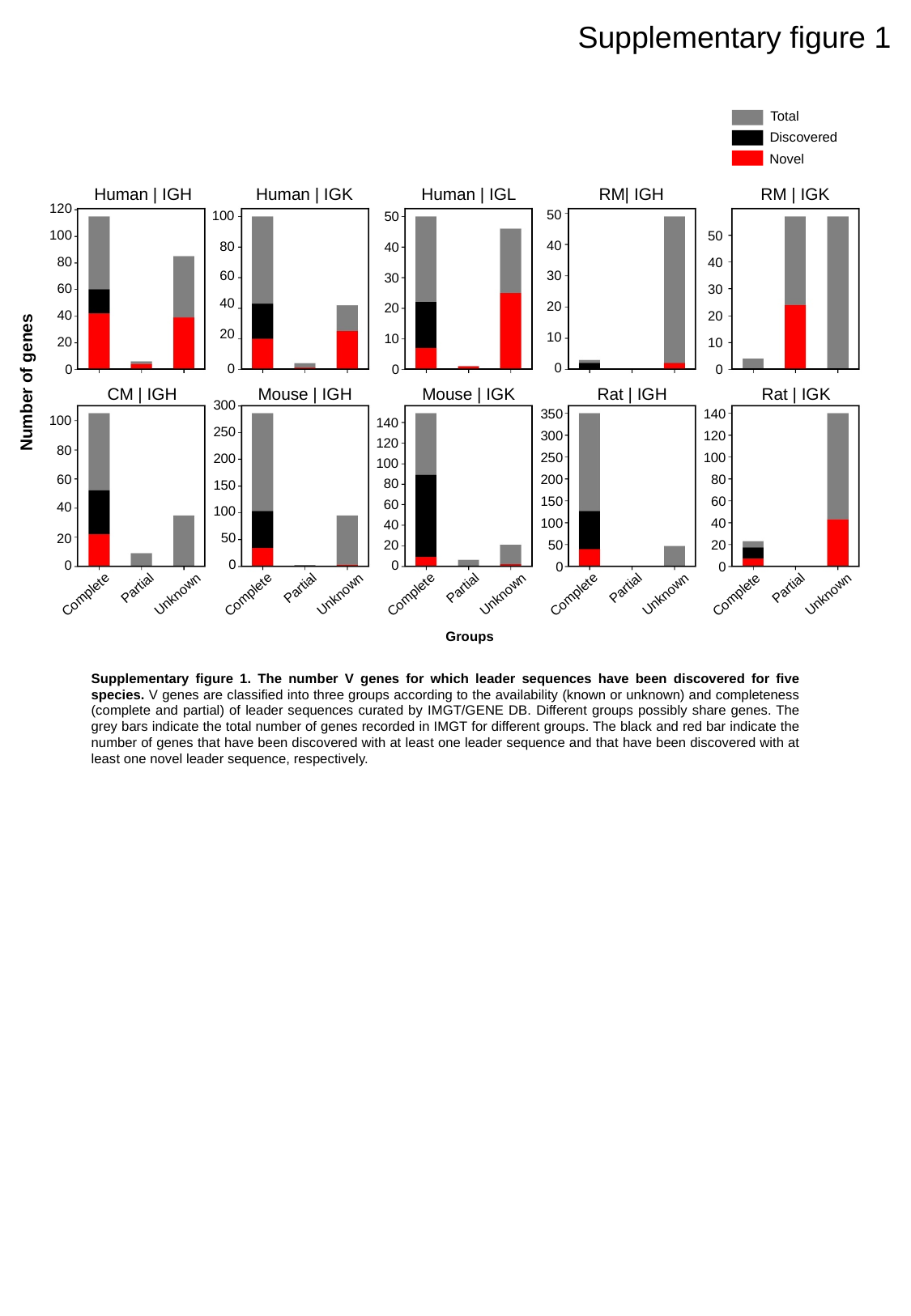

Supplementary figure 1
Total
Discovered
Novel
Human | IGH
Human | IGK
Human | IGL
RM| IGH
RM | IGK
120
100
80
60
40
20
0
50
40
30
20
10
0
100
80
60
40
20
0
50
40
30
20
10
0
50
40
30
20
10
0
Number of genes
CM | IGH
Mouse | IGH
Mouse | IGK
Rat | IGH
Rat | IGK
300
250
200
150
100
50
0
350
300
250
200
150
100
50
0
140
120
100
80
60
40
20
0
100
80
60
40
20
0
140
120
100
80
60
40
20
0
Partial
Unknown
Complete
Partial
Unknown
Complete
Partial
Unknown
Complete
Partial
Unknown
Complete
Partial
Unknown
Complete
Groups
Supplementary figure 1. The number V genes for which leader sequences have been discovered for five species. V genes are classified into three groups according to the availability (known or unknown) and completeness (complete and partial) of leader sequences curated by IMGT/GENE DB. Different groups possibly share genes. The grey bars indicate the total number of genes recorded in IMGT for different groups. The black and red bar indicate the number of genes that have been discovered with at least one leader sequence and that have been discovered with at least one novel leader sequence, respectively.

### Slide 3
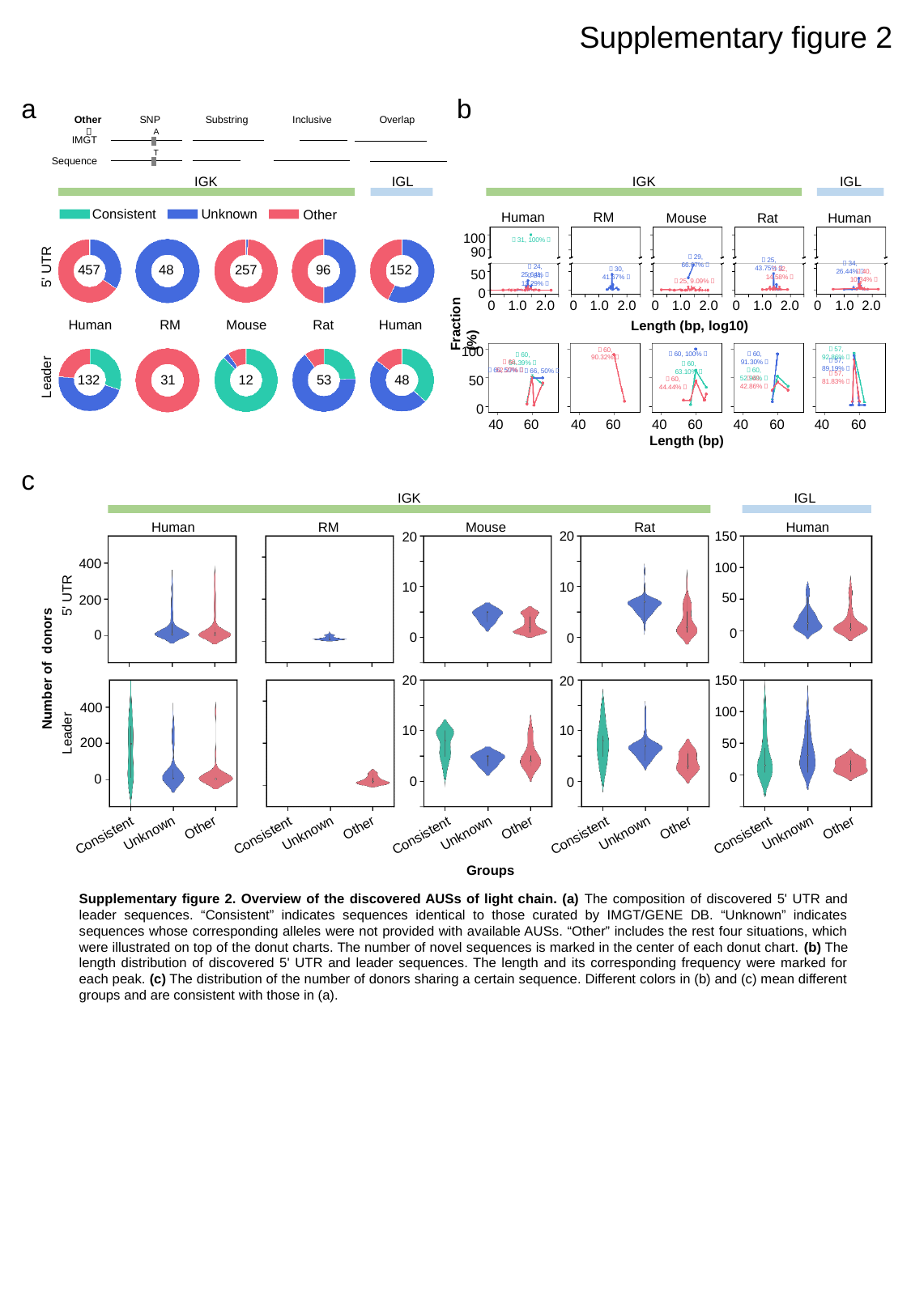

Supplementary figure 2
a
b
SNP
Substring
Inclusive
Overlap
A
IMGT
T
Sequence
Other：
IGK
IGL
IGK
IGL
Consistent
Unknown
Other
Human
RM
Mouse
Rat
Human
100
90
50
0
1.0
2.0
0
1.0
2.0
0
1.0
2.0
0
1.0
2.0
0
1.0
2.0
Length (bp, log10)
0
5' UTR
457
48
257
96
152
（31, 100%）
（29, 66.67%）
（25, 43.75%）
（34, 26.44%）
（24, 25.64%）
（30, 41.67%）
（32, 14.58%）
（40, 10.34%）
（24, 13.29%）
（25, 9.09%）
Fraction (%)
Human
RM
Mouse
Rat
Human
100
50
0
40
60
40
60
40
60
40
60
40
60
Length (bp)
Leader
132
31
12
53
48
（57, 92.86%）
（60, 90.32%）
（60, 91.30%）
（60, 100%）
（60, 54.39%）
（57, 89.19%）
（66, 52.27%）
（60, 63.10%）
（60, 52.94%）
（60, 50%）
（66, 50%）
（57, 81.83%）
（60, 42.86%）
（60, 44.44%）
c
IGK
IGL
Human
RM
Mouse
Rat
Human
150
100
50
0
20
10
0
20
10
0
400
200
0
5' UTR
Number of donors
20
10
0
150
100
50
0
20
10
0
400
200
0
Leader
Other
Unknown
Consistent
Other
Unknown
Consistent
Other
Unknown
Consistent
Other
Unknown
Consistent
Other
Unknown
Consistent
Groups
Supplementary figure 2. Overview of the discovered AUSs of light chain. (a) The composition of discovered 5' UTR and leader sequences. “Consistent” indicates sequences identical to those curated by IMGT/GENE DB. “Unknown” indicates sequences whose corresponding alleles were not provided with available AUSs. “Other” includes the rest four situations, which were illustrated on top of the donut charts. The number of novel sequences is marked in the center of each donut chart. (b) The length distribution of discovered 5' UTR and leader sequences. The length and its corresponding frequency were marked for each peak. (c) The distribution of the number of donors sharing a certain sequence. Different colors in (b) and (c) mean different groups and are consistent with those in (a).

### Slide 4
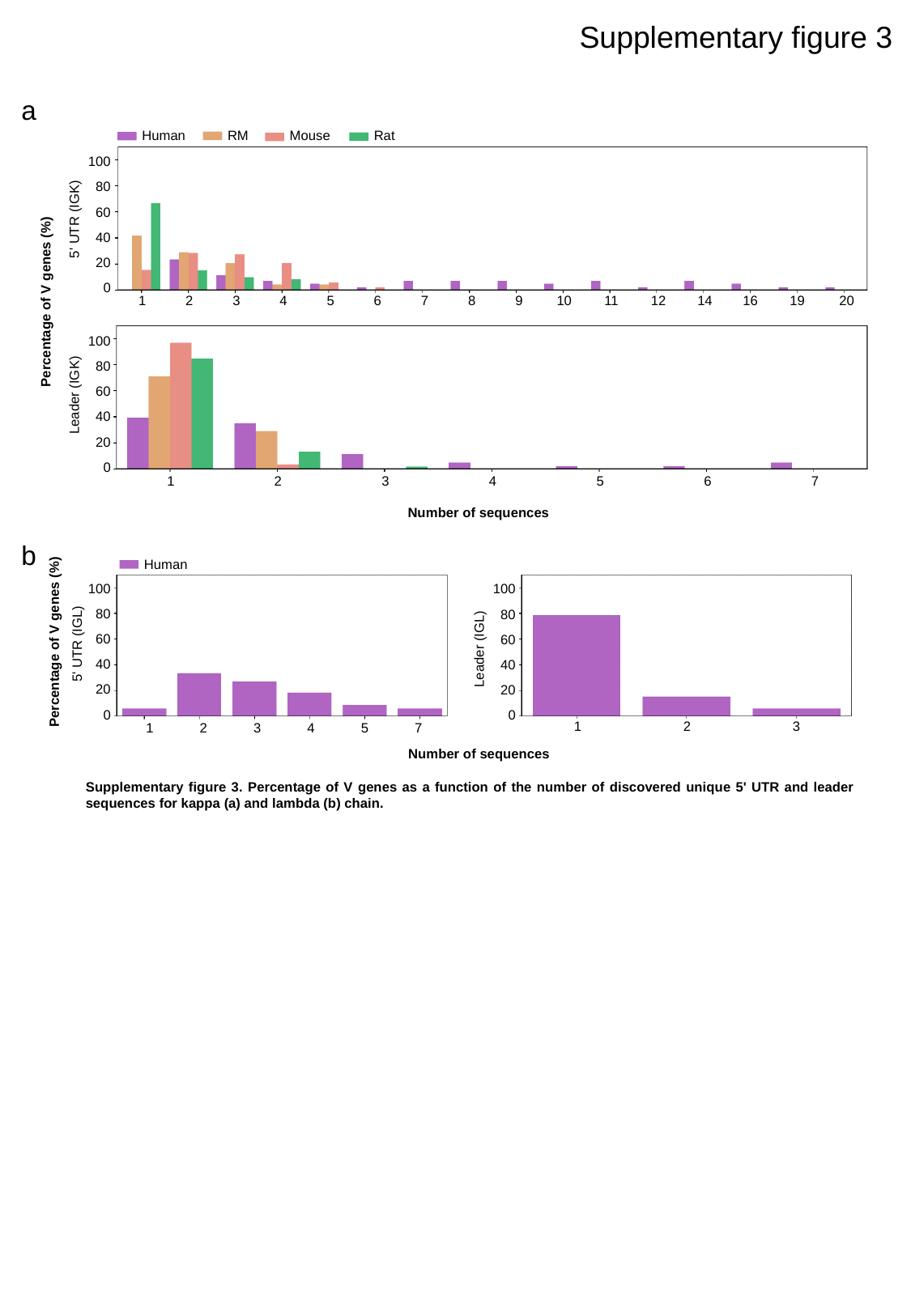

Supplementary figure 3
a
Human
RM
Mouse
Rat
100
80
60
40
20
0
5' UTR (IGK)
1
2
3
4
5
6
7
8
9
10
11
12
14
16
19
20
Percentage of V genes (%)
100
80
60
40
20
0
Leader (IGK)
1
2
3
4
5
6
7
Number of sequences
b
Human
100
80
60
40
20
0
Leader (IGL)
1
2
3
100
80
60
40
20
0
Percentage of V genes (%)
5' UTR (IGL)
1
2
3
4
5
7
Number of sequences
Supplementary figure 3. Percentage of V genes as a function of the number of discovered unique 5' UTR and leader sequences for kappa (a) and lambda (b) chain.

### Slide 5
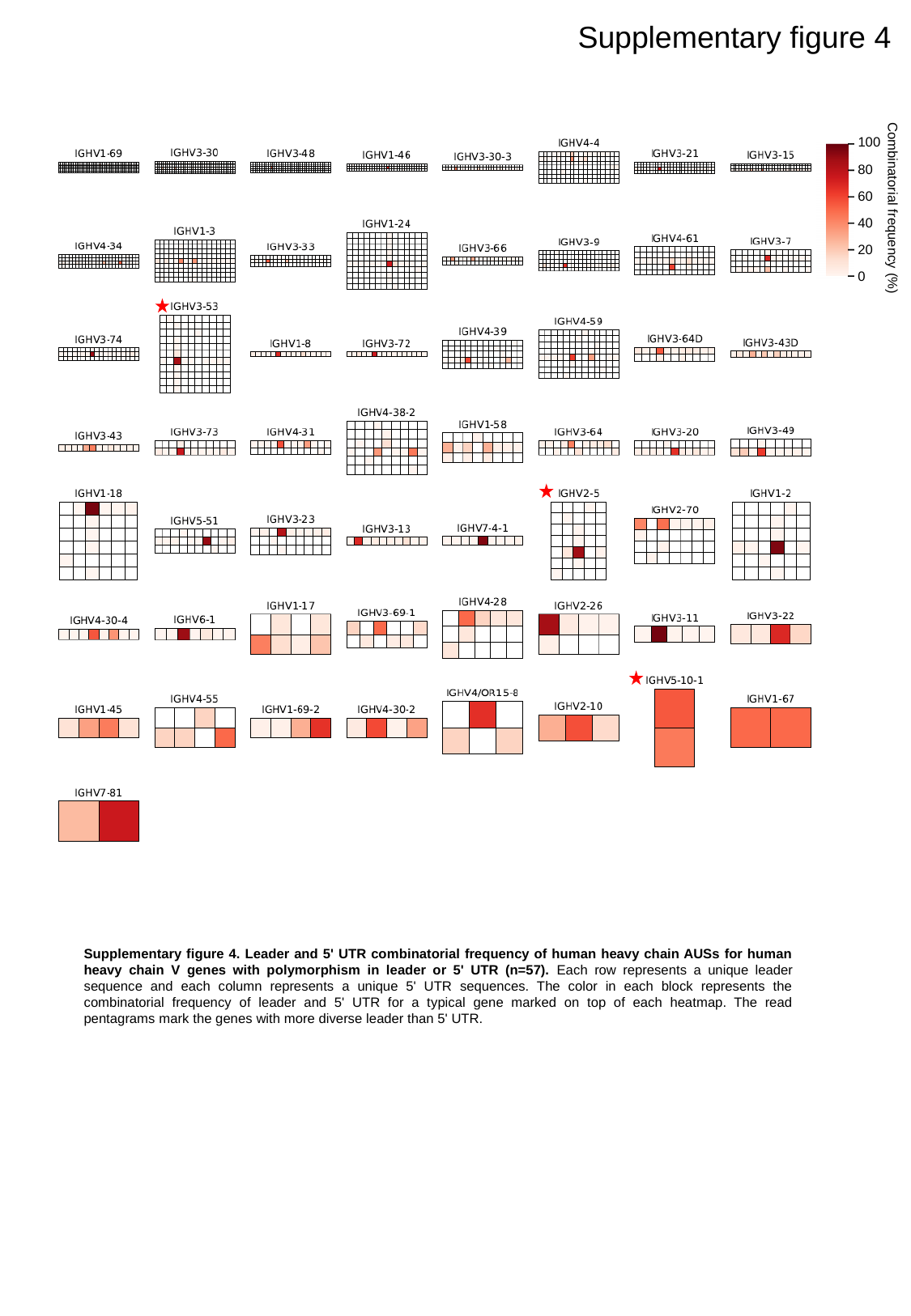

Supplementary figure 4
100
80
60
40
20
0
Combinatorial frequency (%)
Supplementary figure 4. Leader and 5' UTR combinatorial frequency of human heavy chain AUSs for human heavy chain V genes with polymorphism in leader or 5' UTR (n=57). Each row represents a unique leader sequence and each column represents a unique 5' UTR sequences. The color in each block represents the combinatorial frequency of leader and 5' UTR for a typical gene marked on top of each heatmap. The read pentagrams mark the genes with more diverse leader than 5' UTR.

### Slide 6
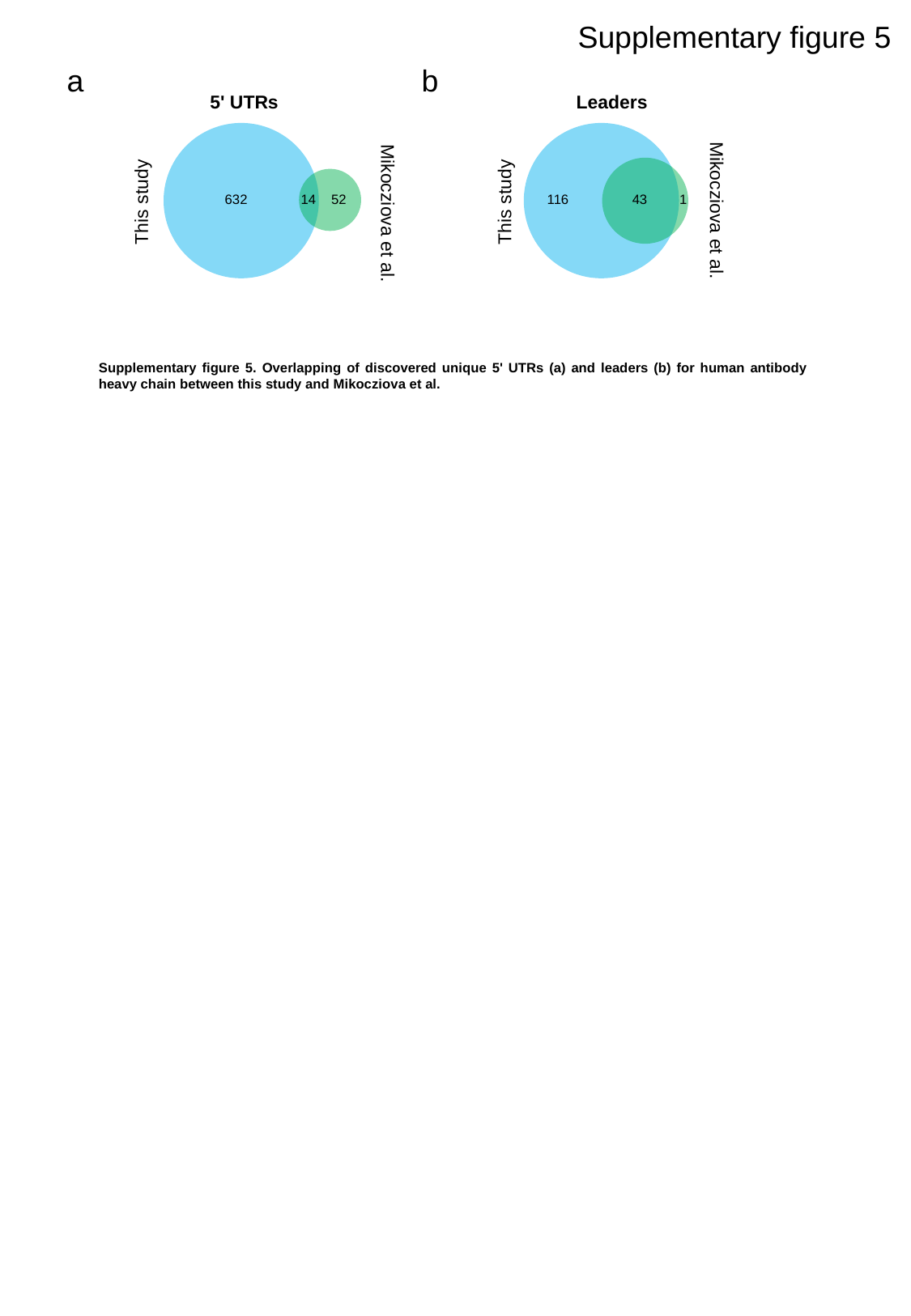

Supplementary figure 5
a
b
5' UTRs
Leaders
This study
This study
632
14
52
116
43
1
Mikocziova et al.
Mikocziova et al.
Supplementary figure 5. Overlapping of discovered unique 5' UTRs (a) and leaders (b) for human antibody heavy chain between this study and Mikocziova et al.

### Slide 7
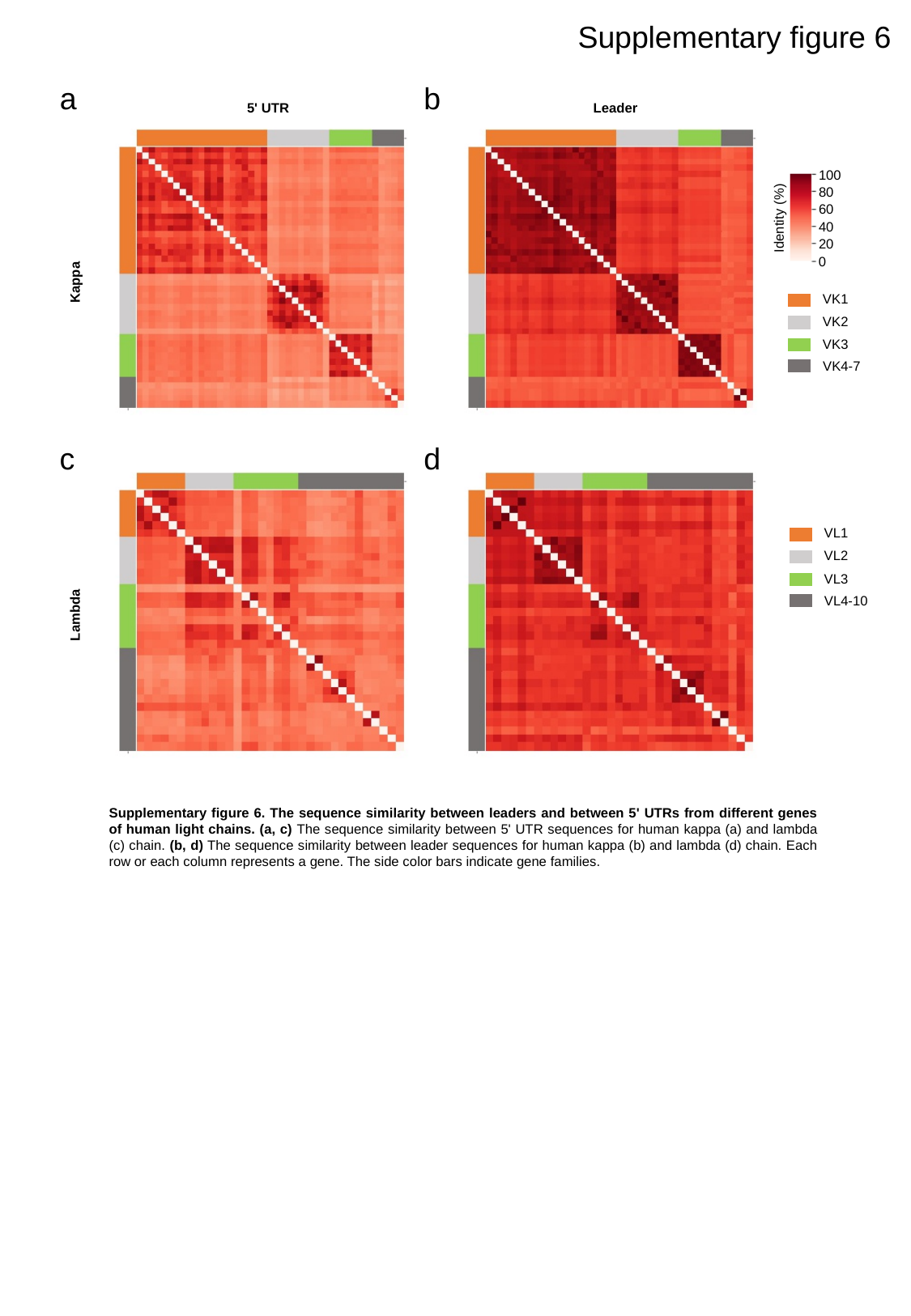

Supplementary figure 6
a
b
5' UTR
Leader
100
80
60
Identity (%)
40
20
0
Kappa
VK1
VK2
VK3
VK4-7
c
d
VL1
VL2
VL3
VL4-10
Lambda
Supplementary figure 6. The sequence similarity between leaders and between 5' UTRs from different genes of human light chains. (a, c) The sequence similarity between 5' UTR sequences for human kappa (a) and lambda (c) chain. (b, d) The sequence similarity between leader sequences for human kappa (b) and lambda (d) chain. Each row or each column represents a gene. The side color bars indicate gene families.

### Slide 8
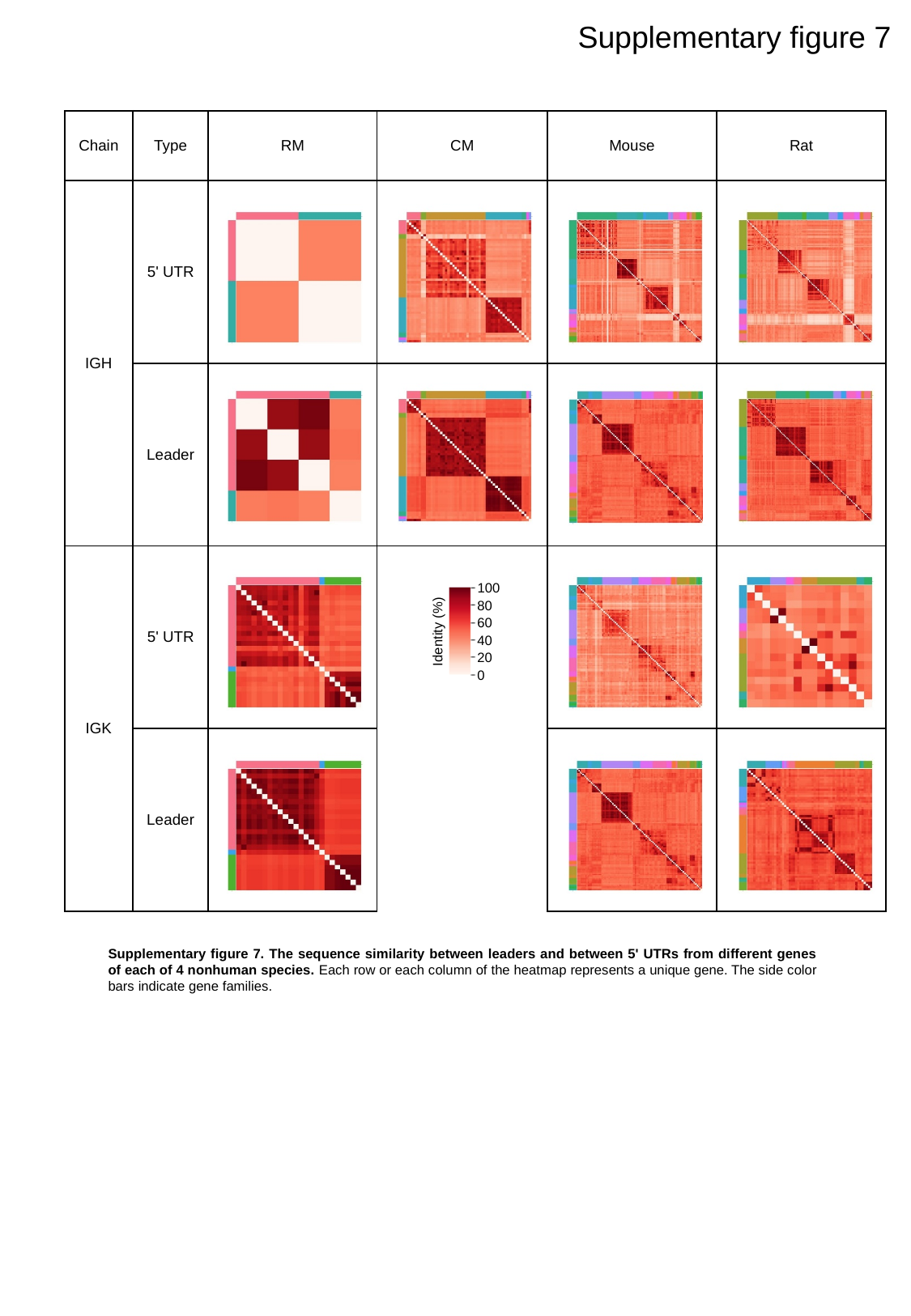

Supplementary figure 7
| Chain | Type | RM | CM | Mouse | Rat |
| --- | --- | --- | --- | --- | --- |
| IGH | 5' UTR | | | | |
| | Leader | | | | |
| IGK | 5' UTR | | | | |
| | Leader | | | | |
100
80
60
Identity (%)
40
20
0
Supplementary figure 7. The sequence similarity between leaders and between 5' UTRs from different genes of each of 4 nonhuman species. Each row or each column of the heatmap represents a unique gene. The side color bars indicate gene families.

### Slide 9
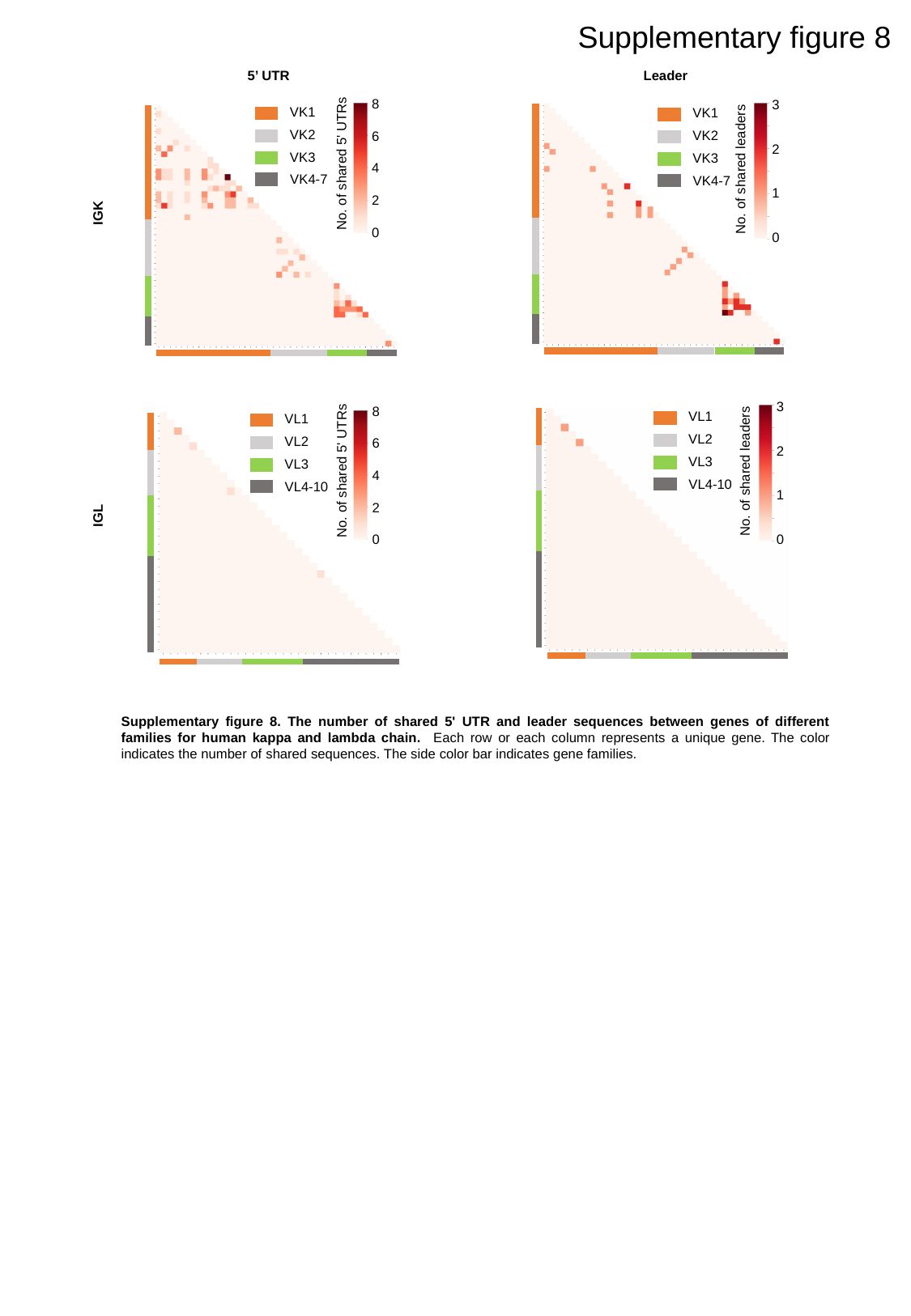

Supplementary figure 8
5’ UTR
Leader
8
6
4
2
0
No. of shared 5' UTRs
3
2
1
0
No. of shared leaders
VK1
VK2
VK3
VK4-7
VK1
VK2
VK3
VK4-7
IGK
8
6
4
2
0
No. of shared 5' UTRs
3
2
1
0
No. of shared leaders
VL1
VL2
VL3
VL4-10
VL1
VL2
VL3
VL4-10
IGL
Supplementary figure 8. The number of shared 5' UTR and leader sequences between genes of different families for human kappa and lambda chain. Each row or each column represents a unique gene. The color indicates the number of shared sequences. The side color bar indicates gene families.

### Slide 10
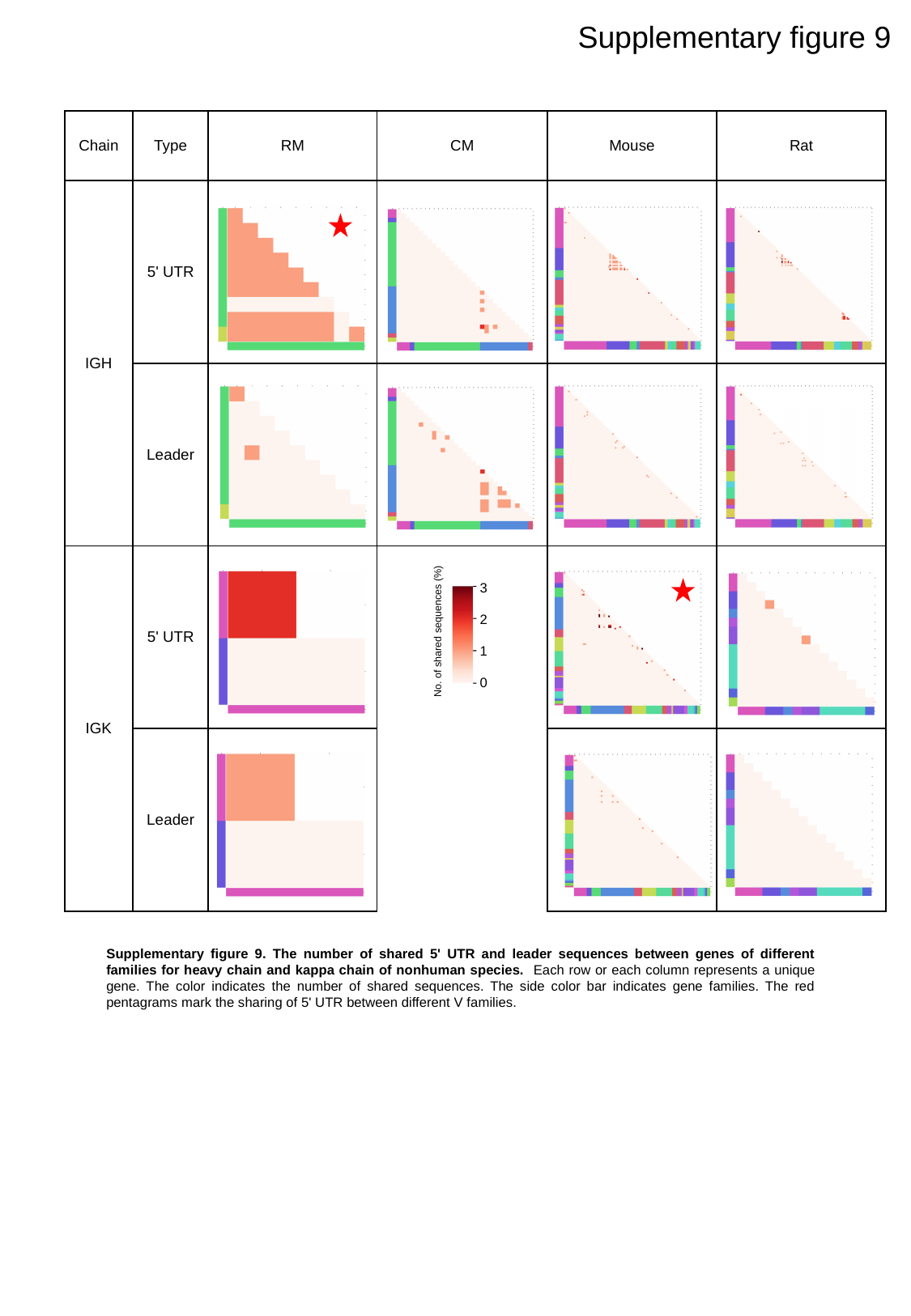

Supplementary figure 9
| Chain | Type | RM | CM | Mouse | Rat |
| --- | --- | --- | --- | --- | --- |
| IGH | 5' UTR | | | | |
| | Leader | | | | |
| IGK | 5' UTR | | | | |
| | Leader | | | | |
3
2
1
0
No. of shared sequences (%)
Supplementary figure 9. The number of shared 5' UTR and leader sequences between genes of different families for heavy chain and kappa chain of nonhuman species. Each row or each column represents a unique gene. The color indicates the number of shared sequences. The side color bar indicates gene families. The red pentagrams mark the sharing of 5' UTR between different V families.

### Slide 11
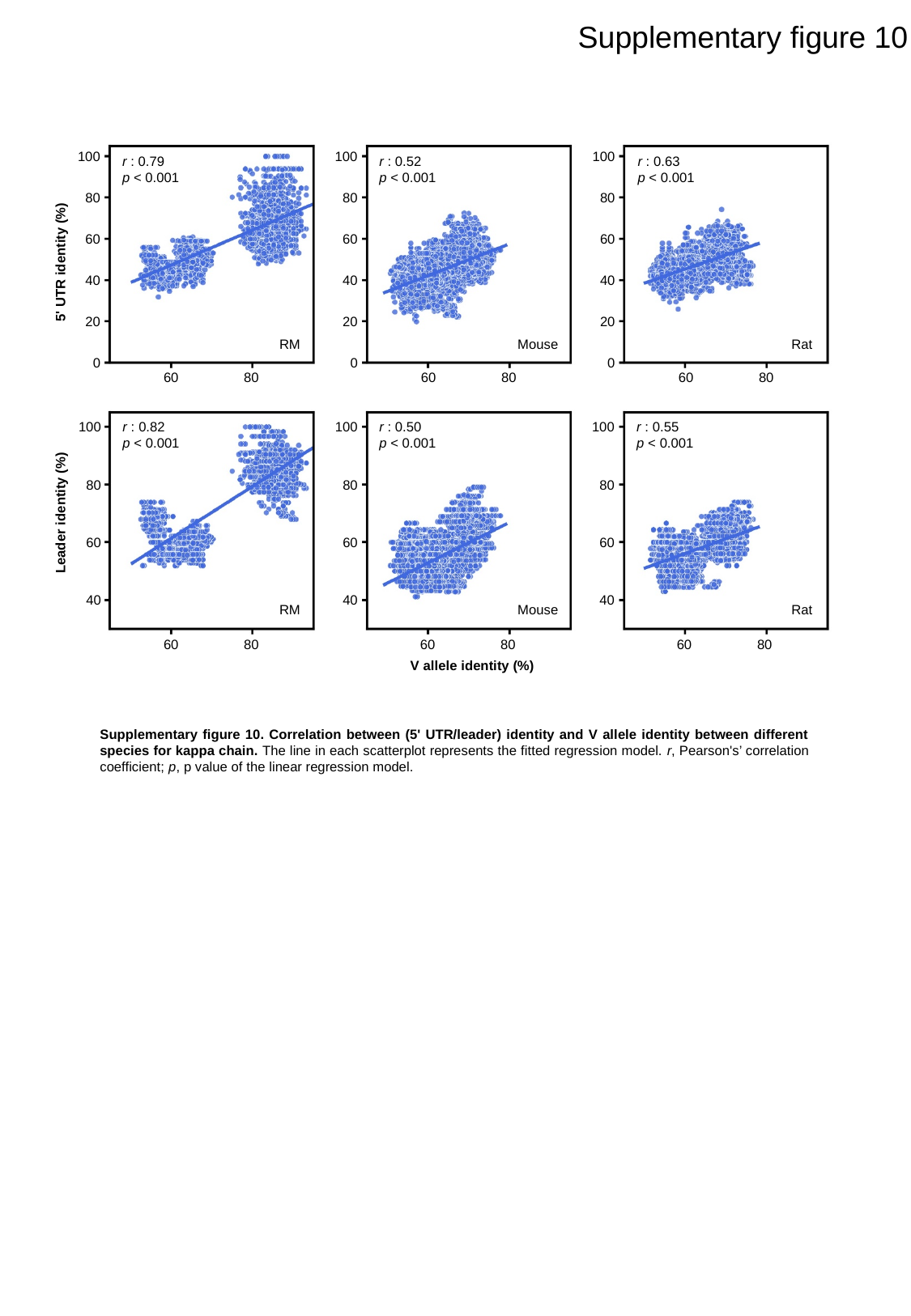

Supplementary figure 10
100
80
60
40
20
0
100
80
60
40
20
0
100
80
60
40
20
0
r : 0.79
p < 0.001
r : 0.52
p < 0.001
r : 0.63
p < 0.001
5' UTR identity (%)
RM
Mouse
Rat
60
80
60
80
60
80
r : 0.82
p < 0.001
r : 0.50
p < 0.001
r : 0.55
p < 0.001
100
80
60
40
100
80
60
40
100
80
60
40
Leader identity (%)
RM
Mouse
Rat
60
80
60
80
60
80
V allele identity (%)
Supplementary figure 10. Correlation between (5' UTR/leader) identity and V allele identity between different species for kappa chain. The line in each scatterplot represents the fitted regression model. r, Pearson's’ correlation coefficient; p, p value of the linear regression model.

### Slide 12
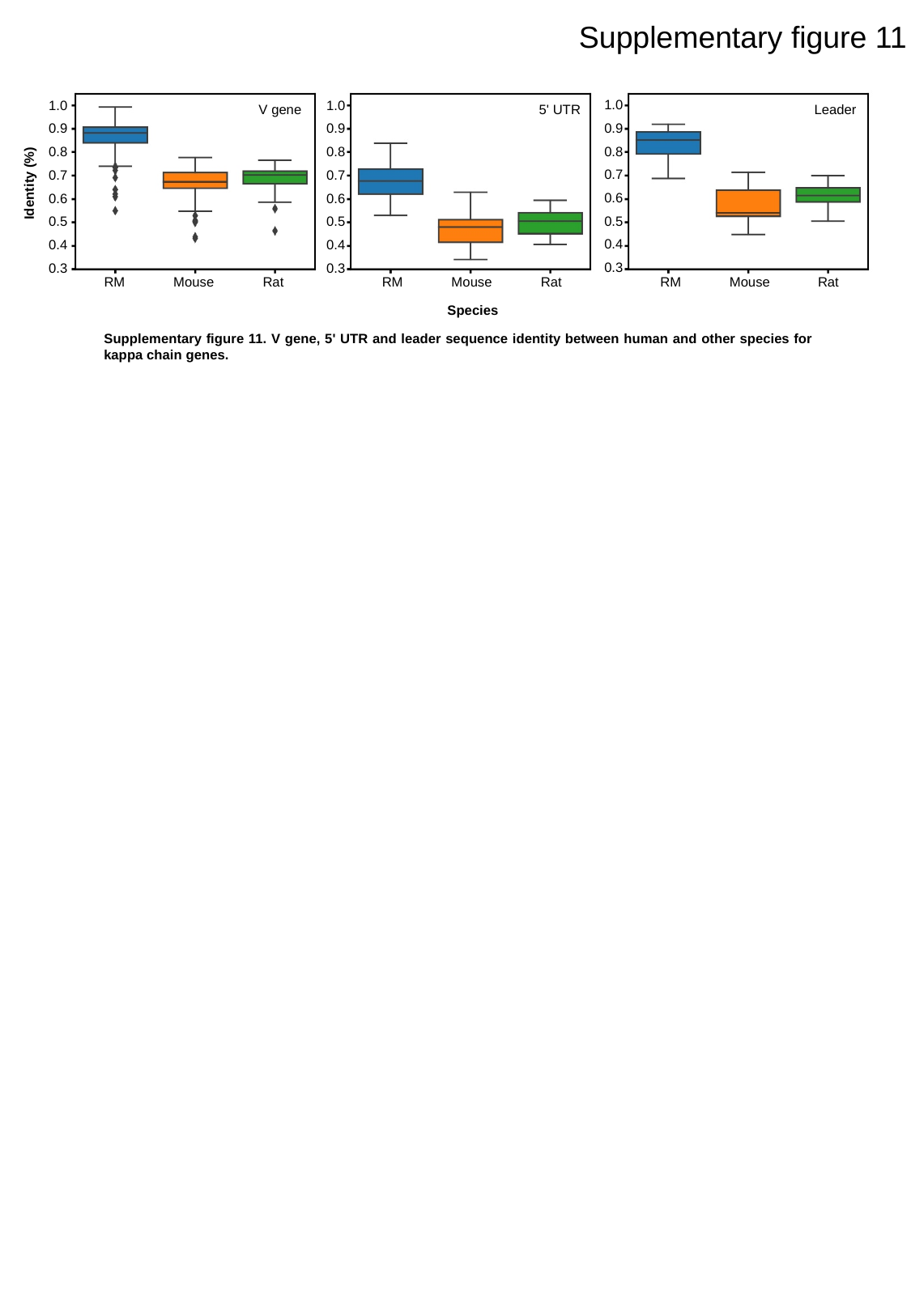

Supplementary figure 11
1.0
0.9
0.8
0.7
0.6
0.5
0.4
0.3
1.0
0.9
0.8
0.7
0.6
0.5
0.4
0.3
1.0
0.9
0.8
0.7
0.6
0.5
0.4
0.3
V gene
5' UTR
Leader
Identity (%)
RM
Mouse
Rat
RM
Mouse
Rat
RM
Mouse
Rat
Species
Supplementary figure 11. V gene, 5' UTR and leader sequence identity between human and other species for kappa chain genes.

### Slide 13
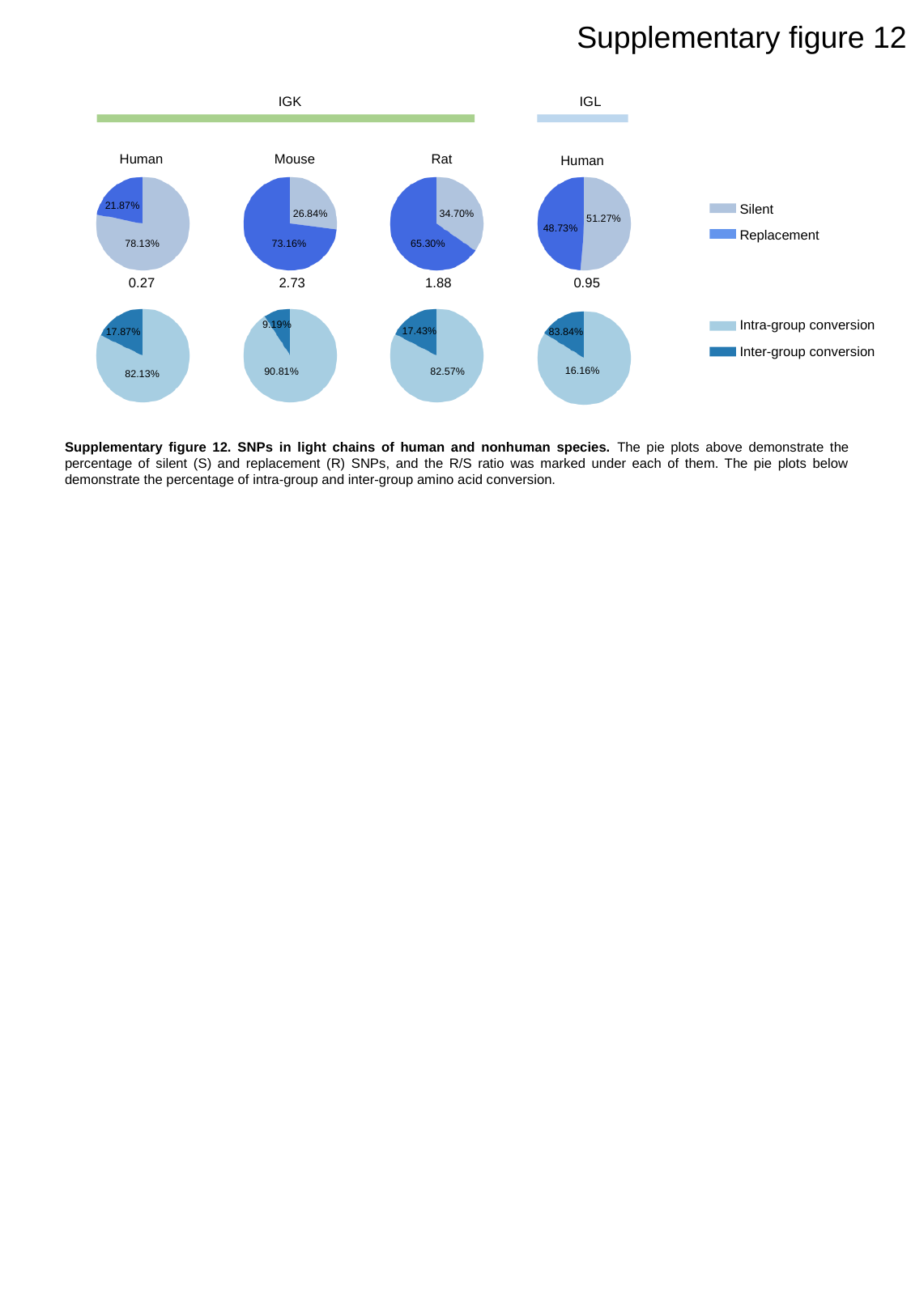

Supplementary figure 12
IGK
IGL
Human
Mouse
Rat
Human
21.87%
Silent
Replacement
26.84%
34.70%
51.27%
48.73%
78.13%
73.16%
65.30%
0.27
2.73
1.88
0.95
Intra-group conversion
Inter-group conversion
9.19%
17.43%
83.84%
17.87%
16.16%
90.81%
82.57%
82.13%
Supplementary figure 12. SNPs in light chains of human and nonhuman species. The pie plots above demonstrate the percentage of silent (S) and replacement (R) SNPs, and the R/S ratio was marked under each of them. The pie plots below demonstrate the percentage of intra-group and inter-group amino acid conversion.

### Slide 14
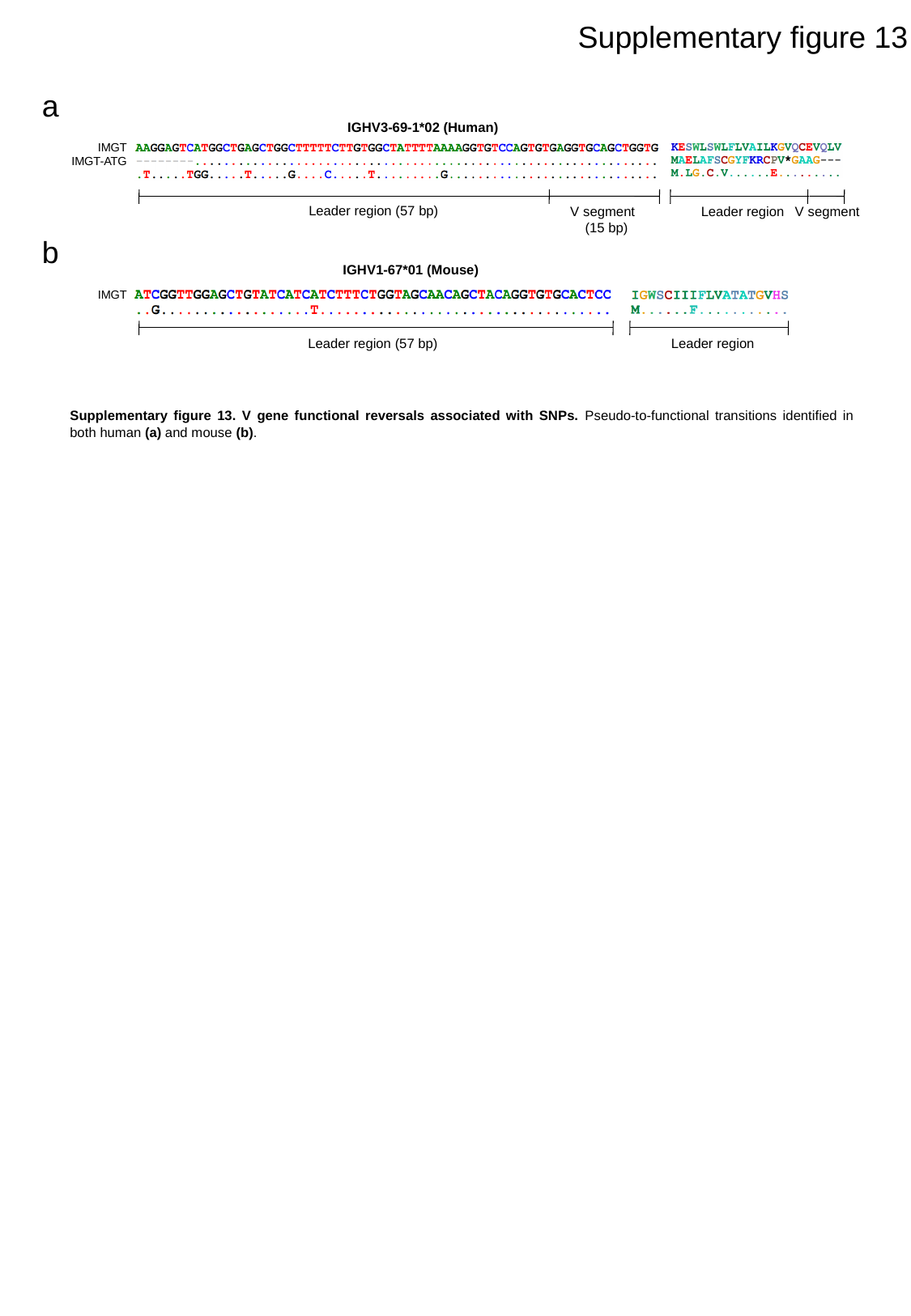

Supplementary figure 13
a
IGHV3-69-1*02 (Human)
IMGT
IMGT-ATG
Leader region (57 bp)
Leader region
V segment
V segment
(15 bp)
b
IGHV1-67*01 (Mouse)
IMGT
Leader region (57 bp)
Leader region
Supplementary figure 13. V gene functional reversals associated with SNPs. Pseudo-to-functional transitions identified in both human (a) and mouse (b).

### Slide 15
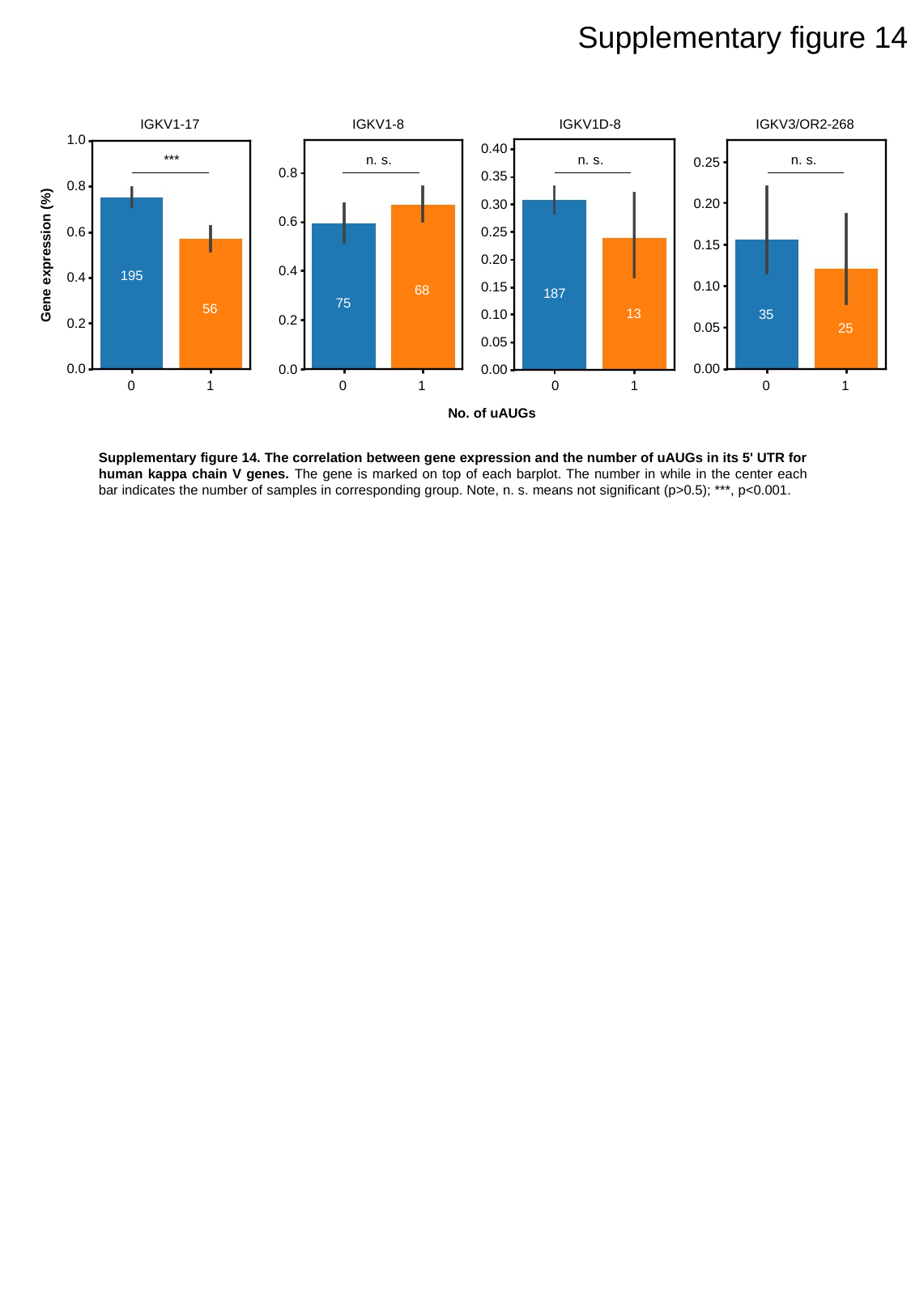

Supplementary figure 14
IGKV1-17
IGKV1-8
IGKV1D-8
IGKV3/OR2-268
1.0
0.8
0.6
0.4
0.2
0.0
0.40
0.35
0.30
0.25
0.20
0.15
0.10
0.05
0.00
***
n. s.
n. s.
n. s.
0.25
0.8
0.6
0.4
0.2
0.0
0.20
0.15
Gene expression (%)
195
0.10
68
187
75
56
13
35
0.05
25
0.00
0
1
0
1
0
1
0
1
No. of uAUGs
Supplementary figure 14. The correlation between gene expression and the number of uAUGs in its 5' UTR for human kappa chain V genes. The gene is marked on top of each barplot. The number in while in the center each bar indicates the number of samples in corresponding group. Note, n. s. means not significant (p>0.5); ***, p<0.001.

### Slide 16
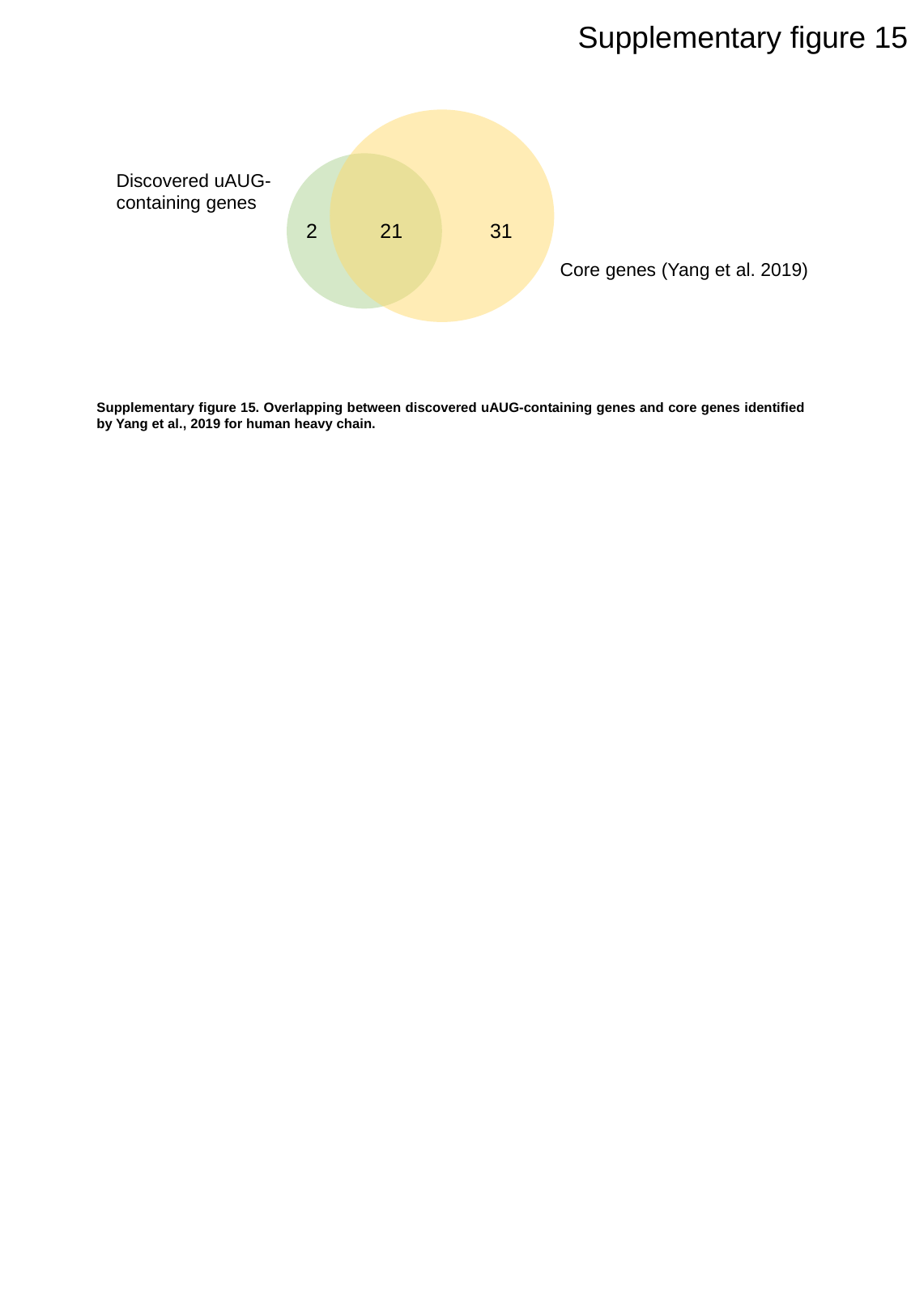

Supplementary figure 15
Discovered uAUG-containing genes
31
2
21
Core genes (Yang et al. 2019)
Supplementary figure 15. Overlapping between discovered uAUG-containing genes and core genes identified by Yang et al., 2019 for human heavy chain.

### Slide 17
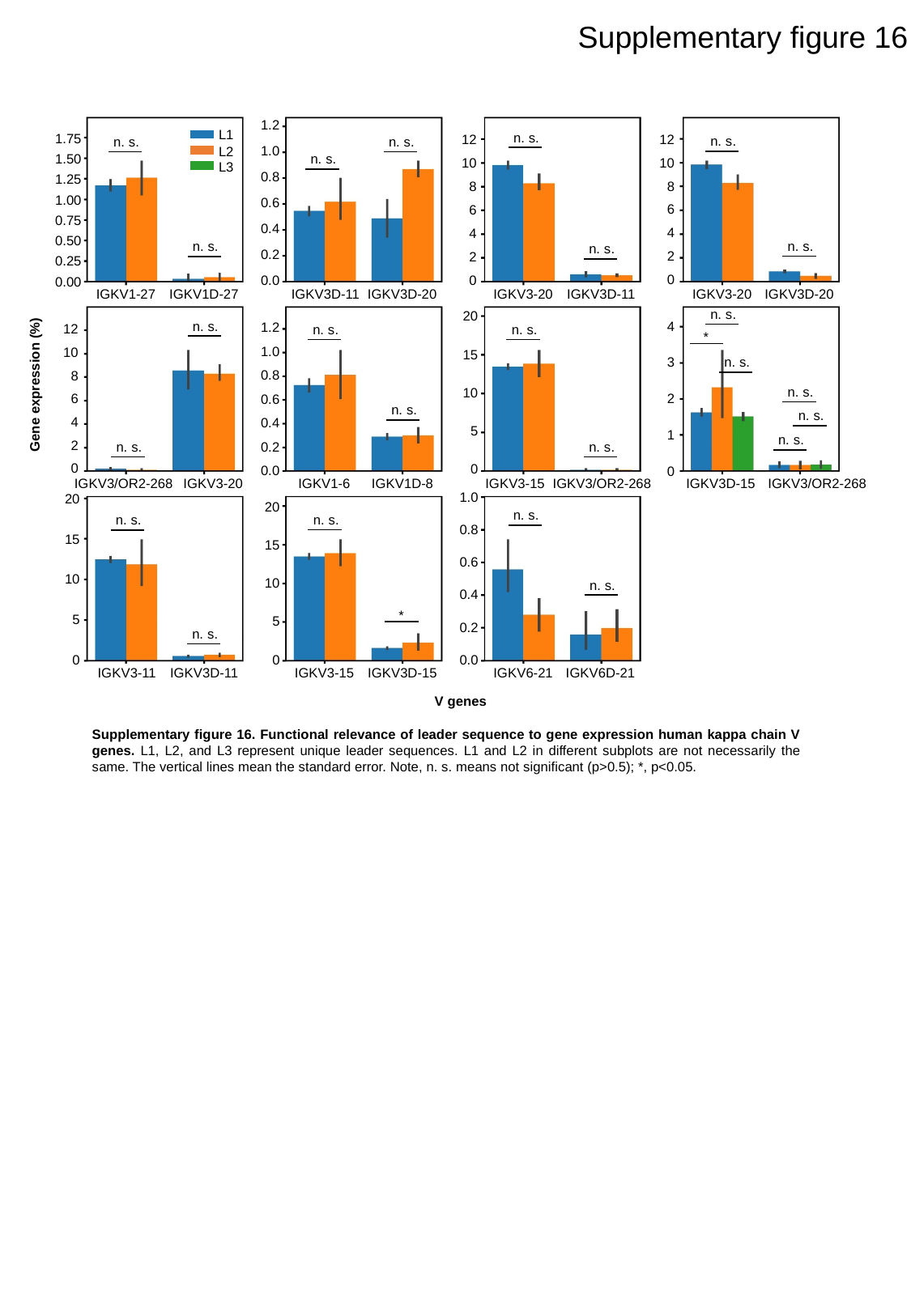

Supplementary figure 16
1.2
1.0
0.8
0.6
0.4
0.2
0.0
L1
n. s.
1.75
1.50
1.25
1.00
0.75
0.50
0.25
0.00
12
10
8
6
4
2
0
12
10
8
6
4
2
0
n. s.
n. s.
n. s.
L2
n. s.
L3
n. s.
n. s.
n. s.
IGKV1-27
IGKV1D-27
IGKV3D-11
IGKV3D-20
IGKV3-20
IGKV3D-11
IGKV3-20
IGKV3D-20
n. s.
20
15
10
5
0
n. s.
4
3
2
1
0
1.2
1.0
0.8
0.6
0.4
0.2
0.0
12
10
8
6
4
2
0
n. s.
n. s.
*
n. s.
Gene expression (%)
n. s.
n. s.
n. s.
n. s.
n. s.
n. s.
IGKV3/OR2-268
IGKV3-20
IGKV1-6
IGKV1D-8
IGKV3-15
IGKV3/OR2-268
IGKV3D-15
IGKV3/OR2-268
1.0
0.8
0.6
0.4
0.2
0.0
20
15
10
5
0
20
15
10
5
0
n. s.
n. s.
n. s.
n. s.
*
n. s.
IGKV3-11
IGKV3D-11
IGKV3-15
IGKV3D-15
IGKV6-21
IGKV6D-21
V genes
Supplementary figure 16. Functional relevance of leader sequence to gene expression human kappa chain V genes. L1, L2, and L3 represent unique leader sequences. L1 and L2 in different subplots are not necessarily the same. The vertical lines mean the standard error. Note, n. s. means not significant (p>0.5); *, p<0.05.
